# Synapses learn to utilize pre-synaptic noise for the prediction of postsynaptic dynamics

**DOI:** 10.1101/2022.04.22.489175

**Authors:** David Kappel, Christian Tetzlaff

## Abstract

Synapses in the brain are highly noisy, which leads to a large trial-by-trial variability. Given how costly synapses are in terms of energy consumption these high levels of noise are surprising. Here we propose that synapses use their noise to represent uncertainties about the activity of the post-synaptic neuron. To show this we utilize the free-energy principle (FEP), a well-established theoretical framework to describe the ability of organisms to self-organize and survive in uncertain environments. This principle provides insights on multiple scales, from high-level behavioral functions such as attention or foraging, to the dynamics of single microcircuits in the brain, suggesting that the FEP can be used to describe all levels of brain function. The synapse-centric account of the FEP that is pursued here, suggests that synapses form an internal model of the somatic membrane dynamics, being updated by a synaptic learning rule that resembles experimentally well-established LTP/LTD mechanisms. This approach entails that a synapse utilizes noisy processes like stochastic synaptic release to also encode its uncertainty about the state of the somatic potential. Although each synapse strives for predicting the somatic dynamics of its neuron, we show that the emergent dynamics of many synapses in a neuronal network resolve different learning problems such as pattern classification or closed-loop control in a dynamic environment. Hereby, synapses coordinate their noise processes to represent and utilize uncertainties on the network level in behaviorally ambiguous situations.

## 1 Introduction

Synapses are inherently unreliable in transmitting their input to the post-synaptic neuron. For example, the probability of neurotransmitter release is typically around 50% [Katz, 1971, Oertner et al., 2002, Jensen et al., 2019] and can be as low as 20% *in vivo* [Borst, 2010]. In other words, up to 80% of synaptic transmissions fail due to release unreliability, providing one of the major sources of noise in the synapse. Pre- and post-synaptic noise sources result in a large trial-by-trial variability in the post-synaptic current (PSC) [Rusakov et al., 2020]. At the same time synapses are very demanding in terms of energy consumption [Pulido and Ryan, 2020], suggesting that a large portion of the body’s energy intake dissipates by the unreliability of synaptic transmission. Similar to biological synapses also neuromorphic technologies are exposed to noise culminating in unreliable synaptic transmission [Indiveri et al., 2013, van De Burgt et al., 2018, Grollier et al., 2020]. The functional implication of noisy synaptic transmission, whether it is a feature or bug in biological and artificial neuronal systems, is therefore highly debated [Maass, 2014, Aitchison et al., 2014, Neftci et al., 2016, Rusakov et al., 2020, Aitchison et al., 2021]. Here, we show that synapses can exploit noisy synaptic transmission to encode their uncertainty about the somatic membrane potential of the postsynaptic neuron. With each synapse doing this, we further show that this enables a neuronal network to encode and utilize uncertainties.

To establish this result we rely on a widely used model framework to describe biological systems that act in uncertain environments: the *free energy principle* (FEP). The FEP is based on the idea that biological systems instantiate an internal model of their environment that allows them to make predictions, take actions and to minimize surprise [Friston, 2010]. A mathematical formulation of surprise can be closely related to the physical notion of free energy, from which the FEP inherits its name. In the FEP formalism an agent uses internal states to form a model of its environment based on perceived stimuli (Fig. 1Ai). In general, these stimuli map only parts of the environment’s true state, implying an unavoidable residual level of uncertainty. To reduce the uncertainty, the agent performs actions to test its predictions about the environment. These actions may lead to new stimuli that provide feedback about the environment’s true state, triggering an update of the internal model. The FEP successfully explains biological mechanisms on various spatial and temporal scales, e.g. dendritic self-organization [Kiebel and Friston, 2011], network-level learning mechanism [Isomura and Friston, 2018], human behavior [Ramstead et al., 2016], and even evolutionary processes [Ramstead et al., 2018].

**Figure 1:**
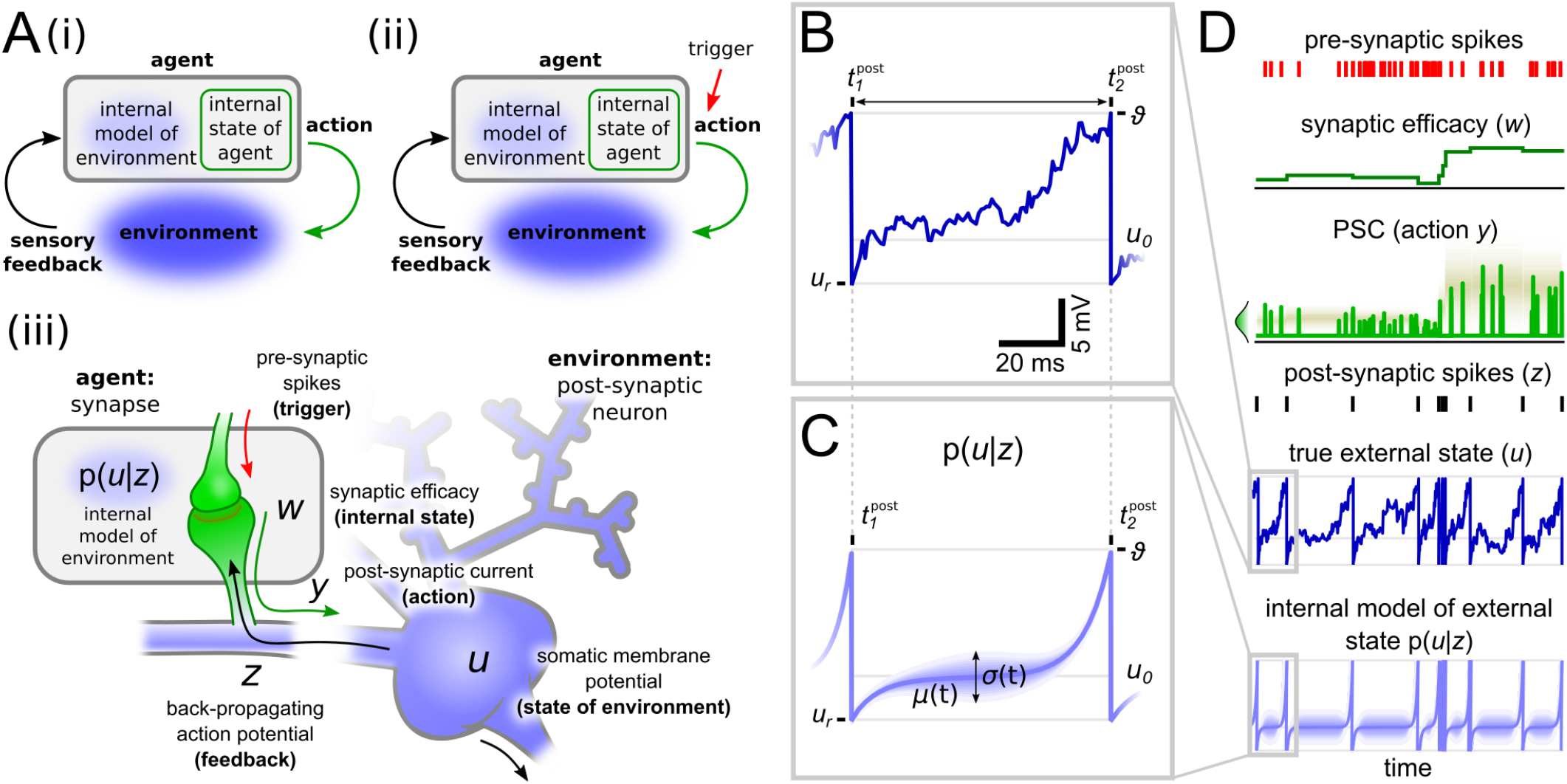
The free energy principle for individual synapses. **A:** i) The FEP considers an agent interacting with the environment through actions. Actions are determined by the agent’s internal state. Sensory feedback from the environment to the agent is used to update the agent’s internal model of the environment. ii) An additional, external trigger can be included into the framework from i) that determines when actions are initialized. iii) The framework shown in ii) can be transferred to a synapse that interacts with its postsynaptic soma. Relevant variables are the synaptic efficacy (internal state), the postsynaptic current (action), the somatic membrane potential (environmental state) of the postsynaptic neuron (environment), and the back-propagating action potential (feedback). **B:** A single trajectory of the somatic membrane potential *u*(*t*) between two action potentials. **C:** The internal synaptic model of the somatic membrane potential can be characterized by the stochastic bridge model providing the probability distribution *p*(*u* | *z*) about the value of *u*(*t*) between two postsynaptic spikes. Solid blue line shows the mean, variance indicated by shaded area. **D:** Illustration of relevant dynamics. Pre-synaptic input spikes (red) trigger synapses to release stochastic postsynaptic currents *y* (light green) with a mean and variance of amplitudes dependent on the synaptic efficacy *w* (dark green). Postsynaptic spike timings reach the synapse through bAPs *z*, constraining the internal model of the somatic membrane potential to the firing threshold *ϑ* and then reset to *u*_*r*_ immediately after every bAP (see panel C). Between bAPs, the internal model estimates the probability density of the membrane potential according to the stochastic process (*µ*(*t*), *σ*(*t*)).

We apply the FEP to individual synapses, arguing that the dynamics of a synapse can be considered as an agent interacting with its cellular environment, and derive a synaptic learning rule by minimizing the free energy of individual synapses. This learning rule enables synapses to adapt their synaptic efficacy to best predict future postsynaptic spiking, which are registered by back-propagating action potentials (bAPs). In contrast to previous approaches (e.g. [Isomura et al., 2016]) that used the FEP to understand the influence of neuromodulatory signals such as Dopamine on synaptic plasticity, we focus here on unraveling the dynamics of a single synapse governed by only locally accessible quantities such as the pre- and postsynaptic-spike times and the current value of the synaptic efficacy. The derivation of the synaptic dynamics relies on a small number of assumptions such that we could solve it in closed from. We call our new model the *synaptic free energy principle* (s-FEP). The emergent synaptic plasticity rule reproduces a number of experimentally observed effects of long-term potentiation (LTP) and depression (LTD) protocols and predicts precise forms for the influence of synaptic and neuron parameters. This result suggests that synapses probe their environment by sending stochastic synaptic currents and integrate the arriving feedback (bAPs) to update their internal state (synaptic efficacy) to better predict the somatic dynamics. Thus, every stochastic release event can be seen as a “*small experiment* “, that is based on previous experience and the outcome of which shapes subsequent future activity. In other words, as we show here, the task of a synapse to learn suitable synaptic responses can be considered as a problem of behaving in a partially unknown environment, where the synaptic noise is being used to properly represent the uncertainty of the synapse about the cellular, environmental state. On the network level, our computer simulations indicate that s-FEP allows several thousand synapses to exploit their synaptic noise to successfully master different learning paradigms despite ambiguous or uncertain inputs.

## 2 Results

After introducing the reasoning to link synaptic properties with the FEP and the fundamentals of the considered model (Section 2.1), we sketch the derivation of the resulting synaptic plasticity rule and compare it with experimental data (Section 2.2). We show that this plasticity rule coordinates the unreliability in synaptic transmission of a group of synapses to drive their joint postsynaptic neuron selectively in a deterministic or probabilistic way (Section 2.3). This successful coordination also functions in feedforward as well as recurrent neuronal networks allowing the system to decode reliably ambiguous stimuli (Section 2.4) or to behave in dynamic environments (Section 2.5).

### 2.1 A synaptic account of the free energy principle

The FEP provides a generic approach to model the behavior of an agent that interacts with its environment. The FEP’s main assumption is that the agent and the environment have physically separated states, that cannot directly influence another. Interaction between the agent’s (*internal*) and the environment’s (*external*) state only takes place through *actions* performed by the agent and *sensory feedback* provided by the environment (see Fig. 1Ai). The FEP suggests a specific method to solve the *internal state* → *action* → *external state* → *feedback* -loop by minimizing the surprise caused by the sensory feedback. This renders an optimization problem that can be solved by maintaining an internal model of the environment, allowing the agent to reason about the true external state and its own uncertainty. Sensory feedback is used to update the internal model and state of the agent such that future actions better help the agent to predict the dynamics of the environment.

The FEP provides a mathematical formalism that can be applied to a wide range of systems of behaving agents and their environments. By employing the FEP on an individual synapse, we find that the interactions between agent and environment are very sparse such that sensory feedback and actions only happen at specific time points. This suggests an event-based view on the FEP where actions (and feedback) are only provided at certain triggering times (Fig. 1Aii). This view allows us to separate the ‘*what* ‘ and ‘*when*’ information flow in the synapse model, which simplifies the analysis compared to previous applications of the FEP.

The intuition behind our synapse-centric s-FEP model is illustrated in Fig. 1Aiii. We consider the synapse as an agent and the postsynaptic neuron as its environment. We consider here as state variable of the neuron the somatic membrane potential *u*, as it determines the neuron’s spiking behavior. However, as suggested by experimental findings [Cornejo et al., 2021], the actual value of the somatic membrane potential is hidden from the synapse and therefore the synapse has to infer the value of *u* from the sparse information that propagates back from the soma into the dendrites. We consider that this sparse feedback is implemented by bAP events *z*, given by the firing times 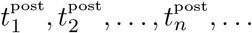 of the postsynaptic neuron, neglecting propagation delays between soma and synapse. Thus, in this model the synapse only receives binary information about the true value of the somatic membrane potential *u*. This information contains whether the somatic membrane potential has recently reached the firing threshold (if 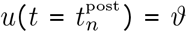, then 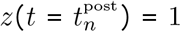 or not (if *u(t)* < *ϑ*, then *z(t)* = 0). As introduced before, in the FEP formalism the feedback causes surprise that is being incorporated into the update of the internal model to better guide the agent’s actions. These interrelations imply in the s-FEP model that bAPs *z* can trigger an update of the synapse yielding new synaptic actions.

The actions of a synapse to interact with the soma are given by the postsynaptic currents (PSCs) *y* (‘*what* ‘) that are released in response to pre-synaptic spikes (‘*when*’). In other words, in the s-FEP the pre-synaptic spikes operate as a trigger for the synapse to initialize an action implemented by PSCs (Fig. 1Aii and Aiii, red arrow). For simplicity we consider the PSC generation as a process of the whole synapse including pre- and postsynaptic mechanisms.Although individual synaptic mechanisms can have specific noise properties [Gontier and Pfister, 2020, Katz, 1971], we integrate pre- and postsynaptic noise sources into one Gaussian noise source that influences the amplitude of PSCs. Thus, at pre-synaptic spike times *t* = *t*^pre^ PSCs are drawn from a general normal distribution with mean and variance being scaled by the synaptic efficacy *w*, in accordance with experimental findings [Meyer et al., 2001].

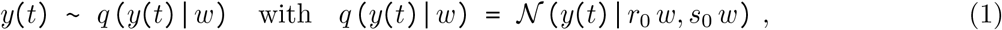

where *r*_0_ > 0 and *s*_0_ > 0 are constants that scale the mean and variance of synaptic currents. At all other times the PSC equals zero (see Fig. 1D). *q(y|w)* determines the distribution over PSC amplitudes for a given synaptic internal state *w*.

To describe the relationship between the feedback and the true external state, the FEP suggests that the agent maintains an internal model of the environment. As the feedback is sparse in time and information (see above), the internal model makes use of a probability distribution over the likelihood of environmental states. This implies that the agent keeps track of its uncertainty about the true external state. In the FEP framework, in general, the environment is too complex to directly infer the external state given the feedback in terms of a posterior probability distribution [Buckley et al., 2017]. However, we find that for the s-FEP we can express this distribution *p* (*u* | *z*) directly in closed form by considering the so-called stochastic bridge model [Corlay, 2013] to approximate the somatic membrane dynamics *u*(*t*) (see Supplementary Text A.2 for more details).

To do so, we have to remind ourselves of the fact that a bAP at time 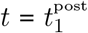 conveys the information to a synapse that the postsynaptic, somatic membrane potential has just reached the firing threshold 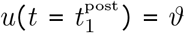. To describe the membrane dynamics between any two bAPs at 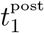 and 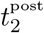, we can utilize that the synaptic uncertainty about the true membrane potential is minimal close to the spike times and maximal between both spikes at 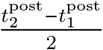. Such an event-based time course of the uncertainty is captured by the stochastic bridge model that determines the distribution of *u*(*t*) given *z* through time-varying mean and variance functions *µ*(*t*) and *σ*(*t*), respectively. Importantly, the shape of *µ*(*t*) and *σ*^2^(*t*) between any pair of postsynaptic spikes depends only on the interspike interval 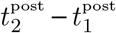. Fig. 1C shows the solution of the stochastic bridge model for 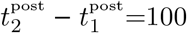 ms that sufficiently captures real membrane dynamics (see Fig. 1B for one example). In other words, we can use the stochastic bridge model to describe the internal representation that the synapse maintains about the somatic membrane dynamics. In the next section we will show that the stochastic bridge model can also be used to infer biologically plausible synaptic plasticity rules to adapt the synaptic efficacies *w*, implying an update of the internal model of the synapse.

### 2.2 Synaptic plasticity as free energy minimization

Learning in the s-FEP means to adapt the synaptic efficacy *w* to minimize the “surprise” caused by bAPs *z*, where surprise is measured with respect to the synapse’s internal model of the soma *p*(*u*|*z*). In other words, the feedback *z* triggers an update of the internal state of the synapse *w*. This update changes the actions of the synapse, namely the PSC amplitudes *y*. The changed actions in turn adapt the dynamics of the somatic membrane potential and, thus, the firing of the postsynaptic neuron that is fed back to the synapse by bAPs *z*. To better understand this loop, we split the effect of the synaptic efficacy into (1) an immediate and (2) a delayed response that are triggered by pre- and post-synaptic firing. Using the event-based view of the s-FEP (Fig. 1Aii), each of these responses can be divided into a ‘*when*’ and a ‘*what* ‘ part. Both responses together determine the adaptation of the synaptic efficacy. The complete process of the synaptic efficacy update Δ*w* is illustrated in Fig. 2A.

**Figure 2:**
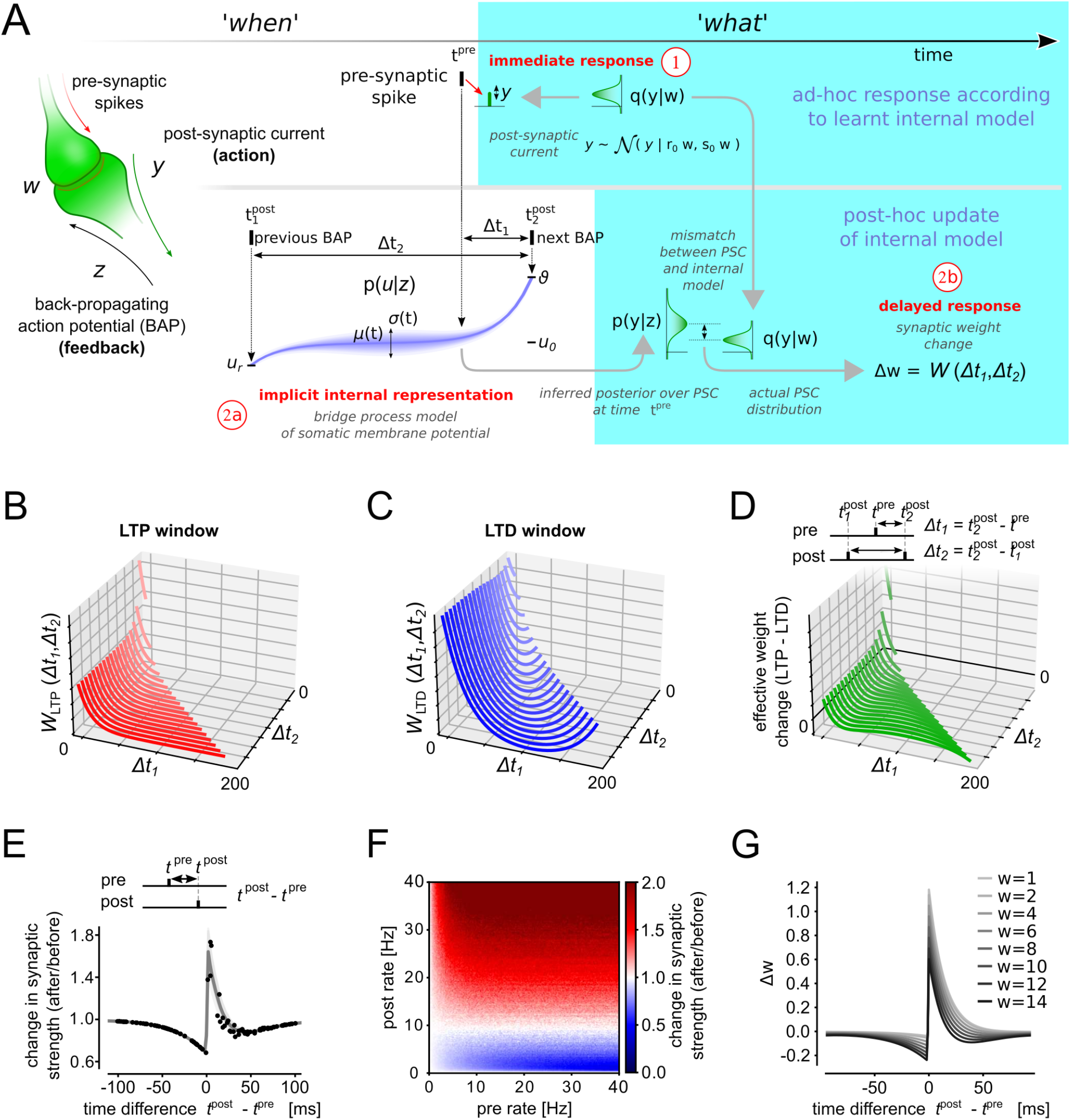
The s-FEP learning rule resembles regulated triplet STDP. **A:** Illustration of the main steps of the s-FEP synaptic learning model. **B**,**C:** The triplet STDP windows *W*_LTP_ (A) and *W*_LTD_ (B) that emerge from s-FEP as a function of the spike timing differences Δ*t*_1_ and Δ*t*_2_. **D:** The effective synaptic efficacy changes that result from the LTP and LTD windows. **E:** Mean synaptic efficacy changes (gray line) and individual trials (black dots) for an STDP pairing protocol. Shaded area indicates variance over trials. **F:** synaptic efficacy changes as a function of pre- and post-rate. **G:** Weight dependence of the s-FEP learning rule for a fixed Δ*t*_2_=200 ms plotted as STDP curve as in (E).

The immediate response (1) determines the action *y* of the synapse. At the time of pre-synaptic firing *t*^pre^ (‘*when*’) a PSC amplitude is generated by drawing a value for *y* from the distribution *q*(*y*|*w*) given in Eq. (1) (‘*what* ‘). The synaptic efficacy *w* determines both, the mean and variance of *y*. The mathematical formulation could be extended comprising two parameters that determine mean and variance separately, but is not investigated here. The immediate response *y*, in other words, constitutes an ad-hoc guess about the firing behavior of the postsynaptic neuron, based on past experience encoded in *w*, before more information is provided by further bAPs (*z*).

The actual update of the synaptic efficacy happens during the delayed response (2). After a further bAP has arrived, the internal model *p*(*u* | *z* is used to update *w* such that the distribution *q*(*y*|*w*) (immediate response) better reflects (or predicts) the postsynaptic firing behavior. The ‘*when*’ part of this update is determined by pre- and post-synaptic firing times. Hereby, the internal model *p*(*u* | *z*) is used to align the relative timing of pre- and post-synaptic firing. This information is then used to estimate the distribution over values of *u* (‘*what* ‘) from which the weight update is inferred. The delayed response thus constitutes a post-hoc correction of the PSC probability distribution *q*(*y*|*w*).

To understand the actual weight update mathematically we have to divide the delayed response (2) into two sub-problems: (2a) We have to invert the internal model *p* (*u* | *z*) to obtain a posterior distribution *p* (*y* | *z*) over synaptic currents *y* that most likely lead to a desired spiking behavior (measured by *z*). (2b) Then we have to reduce the distance between the inferred posterior distribution *p* (*y* | *z*) and the actual distribution over PSCs *q* (*y* | *w*) used in the immediate response (1) to update the synaptic efficacy *w*.

To solve the first sub-problem (2a) we make use of the internal model *p*(*u* | *z*) to directly infer PSCs that are compatible with a given spiking behavior *z*. In Supplementary Text A.3 we show that *p* (*y* | *z*) can be analytically expressed using the stochastic bridge model to describe *p*(*u* | *z*). The resulting solution of the posterior distribution is given by a Gaussian distribution with time-varying mean and variance function *m* and *v*, respectively. At any time *t* = *t*^pre^ the posterior over *y(t)* can thus be written as

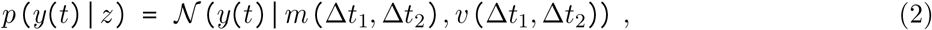

where 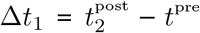 and 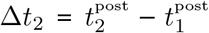 are the relative firing times. This distribution has the property, that it generates PSCs *y* that will, with high probability, result in a spiking behavior *z* when injected into the post-synaptic neuron. Importantly, the functions *m* and *v* only depend on Δ*t*_1_, Δ*t*_2_.

To solve sub-problem (2b), we can use Eqs. (1) and (2), to directly minimize the distance *D(q*| *p*) between *q* (*y* | *w*) and *p* (*y* | *z*). To do so it is sufficient to consider the relative firing times 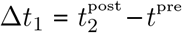 and 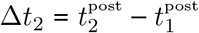 (see Fig. 2A-D). Hereby, Δ*t*_1_ and Δ*t*_2_ can be linked to learning windows of spike-timing-dependent plasticity (STDP, Fig. 2B,C). These learning windows implicitly encode the relevant dynamics of the stochastic bridge model, and thus the internal model does not have to be encoded explicit in every synapse. The synaptic efficacy updates can then be expressed in the form (see Supplementary Text A.4 for a detailed derivation)

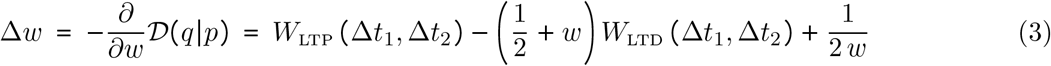

where *W*_LTP_ (Δ*t*_1_, Δ*t*_2_) ≥ 0 and *W*_LTD_ (Δ*t*_1_, Δ*t*_2_) ≥ 0 are triplet STDP learning windows that depend only on the relative timing Δ*t*_1_ and Δ*t*_2_ of pre- and post-synaptic firing, and where *w* denotes the current value of the synaptic efficacy.

In summary we find the main required steps to compute the synaptic weight updates according to the s-FEP model (Fig. 2A). The arrival of a pre-synaptic spike at time *t*^pre^ leads to an ad-hoc response by generating a postsynaptic current *y* according to the internal model *q* (*y* | *w*). When the next bAP arrives at the synapse a post-hoc update of the synaptic efficacy *w* is triggered according to Eq. (3). The probabilistic model Eq. (2) does not have to be explicitly represented in the synapse but is implicit in the shape of the learning windows *W*_LTP_ (Δ*t*_1_, Δ*t*_2_) and *W*_LTD_ (Δ*t*_1_, Δ*t*_2_). Note that the synaptic weight updates strictly follow the separation of the ‘*what* ‘ and ‘*when*’ information flow of event-based FEP (Fig. 1Aii).

The functional form of the two triplet STDP windows is determined by the neuron dynamics (Fig. 1C), and depend on Δ*t*_1_ and Δ*t*_2_ in a nonlinear manner [Pfister and Gerstner, 2006a]. Using Eq. (2) the STDP windows can be expressed as

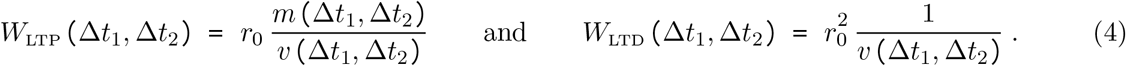

The PSC variance (*v*) has a divisive contribution to both STDP windows. Thus, the learning windows have large values where the uncertainty about *y* is lowest (small values for *v*). In Fig. 2B-D we plot the learning windows for different values of Δ*t*_1_ and Δ*t*_2_. *W*_LTP_ has a potentiating effect which is maximal close to Δ*t*_1_ = 0 (Fig. 2B). This is a manifestation of Hebbian-type learning where close correlations of pre-before post-firing leads to potentiation. *W*_LTD_ is a depression term (Fig. 2C). Both STDP windows show also a strong rate dependence (Δ*t*_2_) as higher firing rates result in less uncertainty about *u t*. Fig. 2D shows the combined effect of the LTP and LTD.

In Fig. 2E we study an STDP pairing protocol where single pre-/post spike pairs with different time lags Δ*t* were presented to a model synapse [Pfister and Gerstner, 2006a]. The resulting synaptic changes with respect to Δ*t* = *t*^post^ − *t*^pre^ closely match experimentally measured STDP windows [Dan and Poo, 2004, Caporale and Dan, 2008]. In Fig. 2F we further study the rate dependence of the s-FEP learning rule. Random pre- and post-synaptic Poisson spike trains were generated with different rates. The learning rule Eq. (3) shows a strong dependence on the post-synaptic firing rate. For low pre- or post-synaptic rates synaptic efficacy changes were zero, moderate post-synaptic rates lead to LTD, whereas high post-synaptic rates manifested in pronounced LTP. This rates-weight-change relation is consistent with previous models of calcium-based plasticity [Graupner and Brunel, 2012].

The learning rule Eq. (3) also includes a dependence on the current synaptic efficacy to regulate the synaptic strength to not grow out of bounds (cf. [Van Rossum et al., 2000, Yger and Gilson, 2015]). The last term becomes effective when synaptic efficacys shrink to values close to zero and prevents the synaptic efficacy from becoming negative (negative weights have no meaning in our model as they also encode variances). The weight dependence of the LTD learning window increases the influence of depression for larger synaptic efficacies. In Fig. 2G we further analyze the weight dependence of the learning rule. STDP protocols for synapses with different initial synaptic efficacies were applied. Small synaptic efficacies (*w*=1 pA) lead to learning windows that are positive for all lags Δ*t* (LTP only). Large synaptic efficacies *w*=12 pA lead to pronounced LTD.

In summary, the s-FEP learning rule contains features of Hebbian learning, STDP and rate-dependent synaptic plasticity to update the synaptic actions (PSC amplitudes) to better predict the somatic membrane dynamics. The learning rule can be best described by a post-pre-post triplet STDP rule [Pfister and Gerstner, 2006a, Pfister and Gerstner, 2006b, Gjorgjieva et al., 2011]. The strength of LTD increases with the synaptic efficacy *w*, which gives rise to a homeostatic effect that prevents synapses from growing out of bounds.

### 2.3 Synapse-level probability matching of firing times

Next, we show how the learned behavior of synapses influences the firing dynamics of the post-synaptic neuron. After the synapse has formed a model of the environment, it can be used to reproduce state trajectories that match the learned behavior. For the s-FEP, this means that synapses install a particular firing pattern *z* through synaptic adaptation. To demonstrate this behavior, we consider here a simple example where a single postsynaptic and many pre-synaptic neurons are repeatedly brought to fire at different fixed offset times (five example pre-synaptic neurons illustrated in Fig. 3A, top). According to the stochastic bridge model, the membrane potential of the postsynaptic neuron evolves according the mean and variance functions illustrated in Fig. 3A, bottom. We forced all neurons to repeatedly fire according to the fixed pattern while the learning rule (3) was active. s-FEP learning installs behavior in the synapses that supports (or predicts) the neuron dynamics (Fig. 3B-D). Individual PSCs after learning are shown for five example synapses in Fig. 3B. The injected currents show high trial-by-trial variability and the amplitude strongly depends on the relative pre- and post-synaptic firing. Despite these variabilities, the summed effect of all PSCs show a clear increasing trend towards the postsynaptic spike (single trial shown in Fig. 3C). In this example, the optimal solutions of synaptic efficacies according to the FEP can be solved analytically. Figure 3D shows the theoretical and simulation results after learning. The synaptic efficacies learn single parameter distributions that encode the theoretically derived *µ*^∗^ and *σ*^∗^. This is further studied in Fig. 3E, where we plot the synaptic efficacies after learning against the euclidean norm 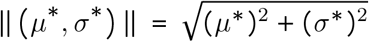. The synaptic efficacies and ‖ (*µ*^∗^, *σ*^∗^) ‖ are strongly correlated (see Supplementary Text A.4 for a theoretical analysis). Fig. 3E shows the estimated mean free energy throughout learning. The free energy steadily declines with learning time.

**Figure 3:**
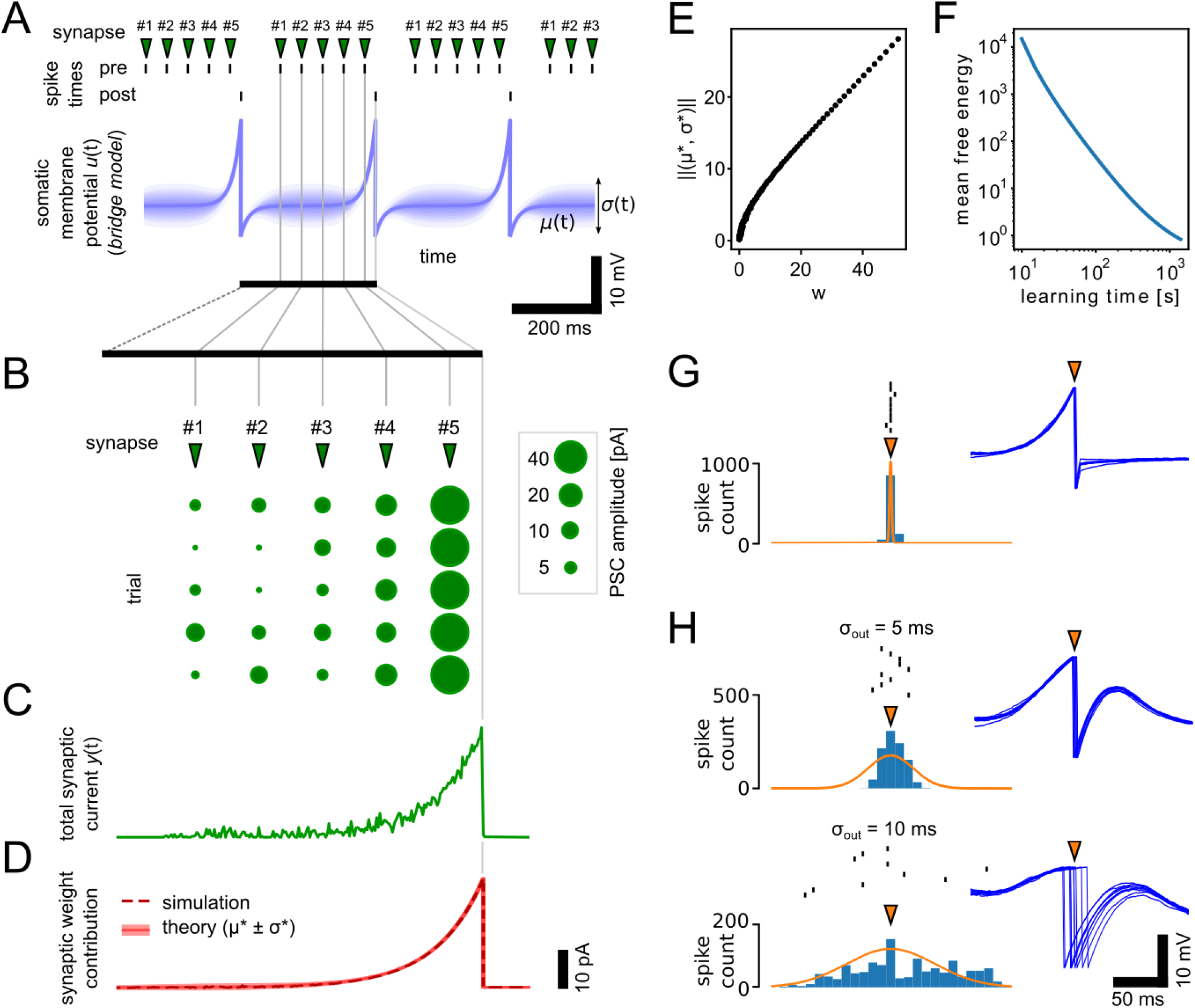
Synapse-level probability matching. **A:** *µ*(*t*) and *σ(t*) of the somatic membrane potential given the stochastic bridge model for a neuron that is brought to fire with a spike interval of 300 ms. Pre-synaptic neurons were brought to fire at fixed time offsets relative to the post-synaptic spikes. **B:** Synapses learn to inject the optimal current that matches the bridge model in (A). Individual current pulses are shown for multiple trials for synapses with different time offsets. **C:** The combined effect of all synapses shown by the summed input current for a single trial. **D:** Synaptic efficacies after learning and weight means and variances predicted by the theory. **E:** Synaptic efficacies after learning are correlated with the euclidean norm of the the theoretically derived *µ*^∗^ and *σ*^∗^ (see panel D). **F:** The mean free energy over all synapses declines throughout learning. **G:** Firing behavior of the neuron after learning when allowed to fire freely in response to input spikes. 10 individual spike times are shown together with histograms over 1000 trials. Insets show membrane dynamics during the 10 trial runs. Orange arrow indicates spike time during learning. **H:** As in (G) but here the output spike times where given by Gaussian distributions of different spreads. Orange arrow indicates here the mean.

Fig. 3F shows the spiking behavior after learning when the post-synaptic neuron was allowed to fire freely in response to the same input spikes that were used during learning. The firing was strongly aligned with the target activity (trial-based variance of firing times was 0.1 ms). Despite their highly stochastic nature (Fig. 3B) synapses have learned to drive the post-synaptic neuron to fire reliably. The synaptic variability can also be exploited to reflect uncertainty in neural firing. We let the postsynaptic neuron learn to fire according to Gaussian distributions of firing times with different spreads *σ*_*out*_ (Fig. 3G, *σ*_*out*_=5 ms and *σ*_*out*_=10 ms). The variability in firing times is reflected in the neuron spiking after learning. During the phase of stochastic firing we observe a high trial-to-trial variability in the dynamics of the membrane potential (insets in Fig. 3G). Note that the pre-synaptic spike times and the LIF neuron model are deterministic here, so the required trial-by-trial variability is generated exclusively by the synapses. Hence, synapses have learned to utilize their intrinsic variability or noise to drive the deterministic neuron to fire according to a defined probability.

### 2.4 Network-level learning using the FEP-derived learning rule

The s-FEP learning rule lends itself to solve supervised and unsupervised learning scenarios on the level of neuronal networks (see Supplementary Text A.5 for a theoretical treatment). To demonstrate this we consider a pattern classification task (Fig. 4A). The network consists of input neurons that project to a set of output neurons. We generated five spike patterns of 200 ms duration (denoted in Fig. 4 by □, ☆, △,◊ and 0) which were used to control the activity of the input neurons.

**Figure 4:**
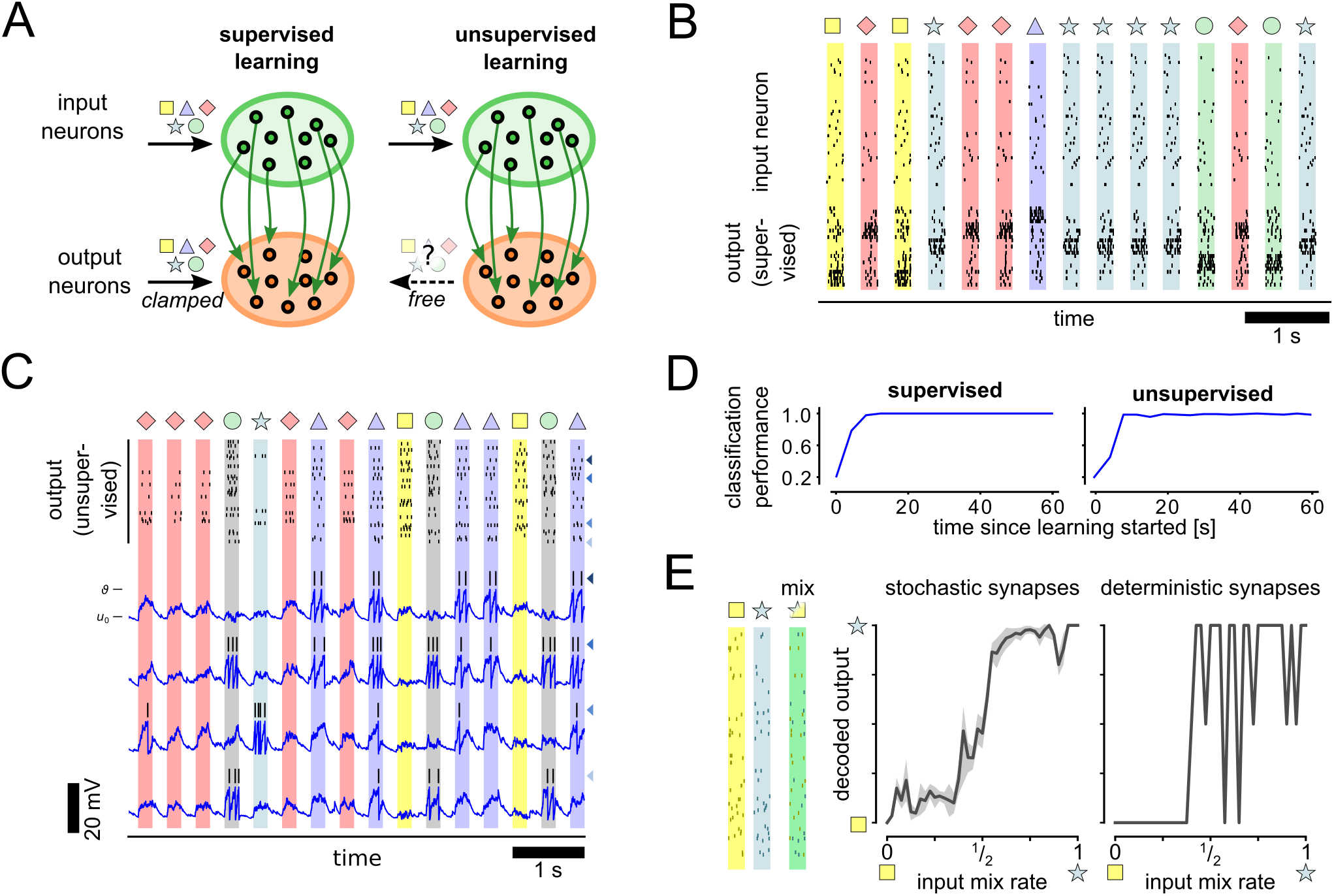
s-FEP learning rule for supervised and unsupervised learning scenarios. **A:** Illustration of the network structure with synapses shown in green being adapted by s-FEP. Five independent spike patterns (□,☆, △,◊, ◯) are presented to the network via the input neurons. Output neurons are either clamped to pattern-specific activity during learning (supervised) or allowed to run freely (unsupervised). **B:** Learning result using the s-FEP rule for the supervised scenario. Typical spiking activity of the network after learning for 60 s. Black ticks show output spike times. **C:** Output activity after learning for the unsupervised scenario. Traces of membrane potentials are shown for selected output neurons (matching color-coded arrows indicate neuron identities). **D:** Classification performance for supervised and unsupervised learning scenario. Classification performance plateaus at near optimal value after about 20 s of learning time for both supervised and unsupervised scenario. **E:** Spike patterns of two input symbols (□,☆) where mixed with different mixing rates (example pattern shows mixing rate 1/2). Uncertainty is reflected in output decoding (left) if inputs are ambiguous (around mixing rate of 1/2). If synapse noise is disabled uncertainty is not represented in the output (right).

Fig. 4B shows the typical network activity after learning for the supervised scenario. The output neurons reliably responded to their preferred pattern. The output neurons had also learned a sparse representation of the input patterns in the unsupervised case (Fig. 4C). Most neurons (46/50) were active during exactly one of the input patterns (e.g. the L:-selective neuron in the top row of Fig. 4C). The remaining neurons showed mixed selectivity and thus got activated by multiple stimulus patterns (see bottom rows of Fig. 4C).

Fig. 4D shows the evolution of the classification performance throughout learning. We used a linear classifier on the network output to recover pattern identities. After learning for 60 s the pattern identity could be recovered by a linear classifier with 100% and 98.8% reliability for the supervised and unsupervised case, respectively (see Fig. 4D). These results demonstrate that the s-FEP learning rule can be applied to supervised learning scenarios and also leads to self-organization of meaningful representations in an unsupervised learning scenario.

To demonstrate the role of noise in the pattern classification task we created ambiguous patterns by mixing the spikes of two patterns (□ and ☆) with different mixing rates (Fig. 4E). Mixing rates of 0 (1) corresponds to a pattern that is identical to □ (☆). This can be encoded by a network with unreliable (left, noisy synapses) and with reliable synaptic transmission (right, without noise). However, intermediate values of the mixing rate result to high levels of ambiguity that are represented in the neural output of the network with unreliable synaptic transmission, but not with noise-free synapses.

### 2.5 Behavioral-level learning using the s-FEP learning in a closed-loop setup

To further investigate the network effects of the s-FEP learning rule, we implemented a closed loop setup where a recurrent spiking neural network controls a behaving agent. So far we have treated the pre-synaptic firing times that trigger the synaptic PSC release as externally given, resulting in the reduced model where synapses only control post-synaptic firing. By considering a recurrent network, the synapse also indirectly gains control over pre-synaptic firing times (see Supplementary Text A.5). The behavioral setup is illustrated in Figure 5A. A fixed goal position *x*_*goal*_ has to be reached starting from *x*_*start*_ in a 3-dimensional task space. The network that was used to learn this task is shown in Fig. 5B. A set of input neurons encoded representations of the current position of the agent’s end effector, which are connected to a the recurrent network of internal state neurons. A set of action neurons were selected from the recurrent network to encode movement directions that are applied to update the agents position. All excitatory synapses in the network were trained using the s-FEP learning rule (3). During training actions are given externally to provide a supervisor signal. The training trajectory (Fig. 5B) had a duration of 1 s. 8 movement trajectories after learning for 220 seconds are shown. Fig. 5C shows corresponding network activity for a single trajectory after learning. The network has learned internal representations to reliably control the agent’s end effector in a closed loop setting.

**Figure 5:**
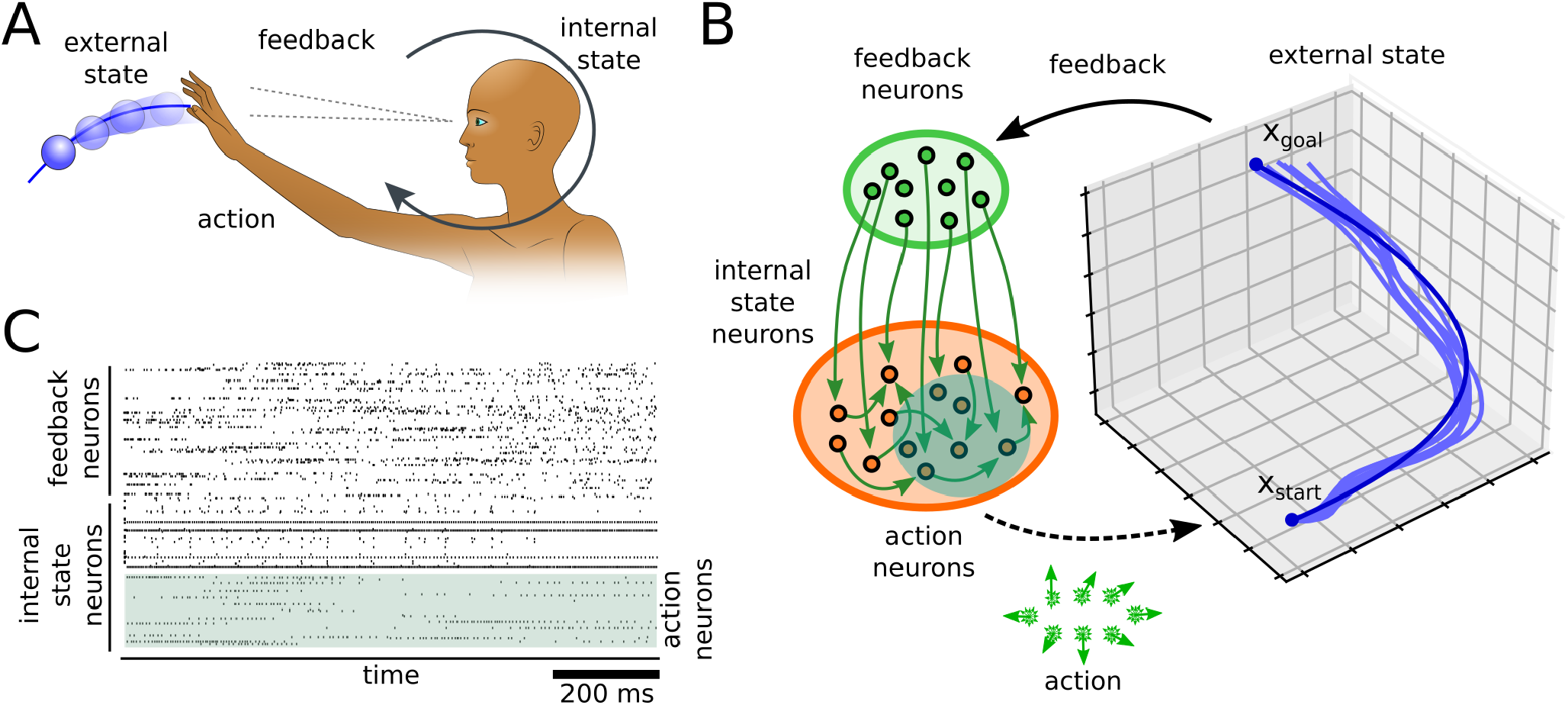
The s-FEP for learning a closed-loop behavior in a recurrent network. **A:** Illustration of the behavior level FEP for an agent that interacts with a dynamic environment. **B:** A spiking neural network interacting with an environment using s-FEP to learn a control policy. The activity of action neurons controls the movement of an agent in a 3-dimensional environment. Feedback about the position of the agent is provided through feedback neurons. The policy to navigate the agent is learned through s-FEP between feedback and action neurons. Training trajectory (dark blue) and 8 spontaneous movement trajectories generated by the network after learning (light blue) are shown. **C:** Spike train generated spontaneously by the network after learning corresponding to one movement trajectory in (B).

## 3 Discussion

The FEP has been praised for its ability to describe biological phenomena on different spatial and temporal scales [Friston, 2010,Friston, 2012]. Here we started from the lower end of the spatial hierarchy: individual synapses that learn to interact with their postsynaptic neuron. To the best of our knowledge, this is the first time that the FEP is applied to the subcellular level to derive local learning rules, proposing a new role of pre-synaptic noise. Previous models suggest that pre-synaptic noise increases the energy and information transmission efficiency of synapses [Neftci et al., 2016,Schug et al., 2021,Levy and Baxter, 1996,Levy and Baxter, 2002,Harris et al., 2012]. We show that pre-synaptic noise can also be utilized to encode uncertainty about the somatic membrane potential, providing a theory on the role of noise on the postsynaptic side. We also demonstrate a first step investigating the functional implication of s-FEP on the network and behavioral level. This complements previous results to derive detailed properties of neural networks based on the FEP [Palacios et al., 2019, Isomura and Friston, 2020].

### 3.1 Prior related work

The FEP and the related theory of predictive coding have been very successful in explaining animal behavior and brain function [Rao and Ballard, 1999, Friston, 2005, Friston, 2010, Chalk et al., 2018]. On the neuron and network level, previous models utilized the FEP to derive learning rules for reward-based learning and models of the dopaminergic system [Friston et al., 2014]. [Isomura et al., 2016] used the FEP to derive synaptic weight updates with third factor modulation using dopamine-like signals. A number of previous studies have approached the problem of deriving learning rules from related variational Bayesian inference theory [Deneve, 2008, Brea et al., 2013, Rezende and Gerstner, 2014, Jimenez Rezende et al., 2011, Rezende et al., 2014] and information-theoretic measures [Toyoizumi et al., 2005, Buesing and Maass, 2008, Buesing and Maass, 2010, Linsker, 1988]. In [Urbanczik and Senn, 2014] a model for dendritic prediction of somatic spiking was proposed. Different to the s-FEP approach, the uncertainty about the somatic membrane potential was not represented in these models.

Recently it was shown that the FEP may also provide an interesting alternative to the error back-propagation algorithm for learning in deep neural networks [Whittington and Bogacz, 2017, Millidge et al., 2020]. The s-FEP complements these results with a bottom-up approach for spiking networks. An important property of the s-FEP learning rule is that synaptic updates only depend on the timing of pre- and post-synaptic spikes, which makes the model well suited for event-based neural simulation [Pecevski et al., 2014, Peyser et al., 2017] and new brain-inspired hardware [Mayr et al., 2019, Davies et al., 2018]. Therefore, s-FEP learning is a promising candidate to control unreliable signal transmission in diverse neuromorphic technologies [Indiveri et al., 2013, van De Burgt et al., 2018, Grollier et al., 2020] and even to exploit the unreliability for learning.

### 3.2 Testable experimental predictions

Direct experimental evidence for the FEP in cultured neurons was provided by [Isomura et al., 2015]. In [Isomura and Friston, 2018] a FEP-based encoding model was formulated and could account for observed electrophysiological responses in vitro. Evidence for predictive coding is also abundantly available in *in vivo* recordings of neural activity and brain anatomy [Bastos et al., 2012, Kanai et al., 2015, Barascud et al., 2016, Driscoll et al., 2017, Kostadinov et al., 2019]. However, the FEP has been often criticized for being too general to make any falsifiable experimental predictions [Friston et al., 2012]. In contrast, the s-FEP proposed here makes very precise predictions about the interplay of synaptic and neural dynamics. Our model directly predicts that synapses should be stochastic to effectively encode uncertainties about the somatic membrane potential. Furthermore, the s-FEP makes precise predictions about the synaptic plasticity changes in post-pre-post spike pairing protocols and the dependence on the synaptic weight (Fig. 2).

### 3.3 Conclusion

In summary, we have presented a synapse-centric account of the FEP that views synapses as agents that interact with their post-synaptic neuron much like an organism interacts with its environment. The emerging s-FEP learning rule matches experimentally observed synaptic mechanisms at a high level of detail. Our results complement previous applications of the FEP on the system and network level and demonstrates that manifestations of the FEP can be identified even on the smallest scales of brain function. In contrast to this prior work our model synapses use only local information and yields triplet STDP dynamics which can be directly tested against experiments. The emergent learning algorithm is fully event-based, i.e. computation only takes place when pre- and post-synaptic spikes arrive at the synapses. The model is therefore very well suited for event-based neural simulation and brain-inspired hardware.

## Methods

### Neuron model

We used the leaky integrate and fire (LIF) neuron model [Gerstner et al., 2014] in all experiments, where the somatic membrane potential *u*(*t*) at time *t* > 0 follows the dynamics

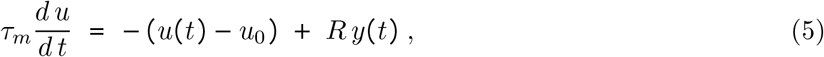

where *τ*_*m*_ is the membrane time constant, *u*_0_ is the resting potential and *R* the membrane resistance. *y*(*t*) is the external input current into the neuron and denotes the effect of afferent synaptic input at time *t*. When the membrane potential reaches the threshold *ϑ*, the neuron emits an action potential, such that the spike times *t*_*f*_ are defined as the time points for which the criterion

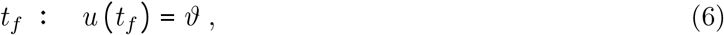

applies. Immediately after each spike the membrane potential is reset to the reset potential *u*_*r*_ [Gerstner et al., 2014]

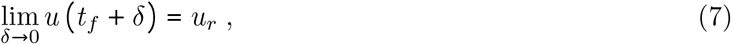

and we define the initial state of the neuron *u*(0) = *u*_*r*_.

In Fig. 4C and Fig. 5 we used a simple threshold adaptation mechanism to control the output rate of the neurons. Individual firing thresholds *ϑ* where used for every neuron. Thresholds were decreased by a value of 10^−5^ mV in every millisecond and increased by 10^−3^ mV after every output spike. Thresholds values were clipped from below at the resting potential *u*_0_.

### Synapse model and Learning rule

A detailed derivation of the synapse model can be found in the Supplementary Text. In Supplementary Text A.1 we summarize the main features and assumptions underlying the s-FEP. In Supplementary Text A.2 we derive the internal model *p* (*u* | *z*). In Supplementary Text A.3 we define the PSC distribution *q* (*y* | *w*). In Supplementary Text A.4 we derive the synaptic efficacy updates (3). In Supplementary Text A.5 we develop the network-level s-FEP model.

We use a stochastic synapse model of input-dependent PSCs, where the variability is proportional to the synaptic efficacy *w* [Yang and Xu-Friedman, 2013]. The parameter *s*_0_ of the stochastic PSC model (1) was chosen to be the Gaussian approximation of the Binomial distribution, with *s*_0_ =*r*_0_ (1− *r*_0_) On a pre-synaptic spike a current pulse with amplitude *y* drawn from a Gaussian distribution *𝒩* (*y* | *r*_0_ *w, r*_0_ (1 − *r*_0_) *w*), with scaling constant *r*_0_, is injected into the post-synaptic neuron. Otherwise_e_ the synaptic inputs *y*(*t*) were 0.

If not stated otherwise the synaptic efficacies *w* were updated using the learning rule Eq. (3), where *W*_LTP_ (Δ*t*_1_, Δ*t*_2_) and *W*_LTD_ (Δ*t*_1_, Δ*t*_2_) are the triplet STDP windows as depicted in Fig. 2. In Supplementary Text A.4 we show in detail that the synaptic efficacy updates (3) minimize the free energy *F* (*z, w*) of the back-propagating action potentials *z* with respect to the synaptic efficacy *w*. We further show that the triplet STDP windows can be defined analytically using the stochastic bridge model with time varying mean and variance functions *µ* (Δ*t*_1_, Δ*t*_2_) and *σ*^2^ (Δ*t*_1_, Δ*t*_2_), respectively

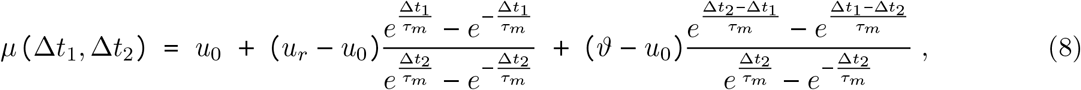

and

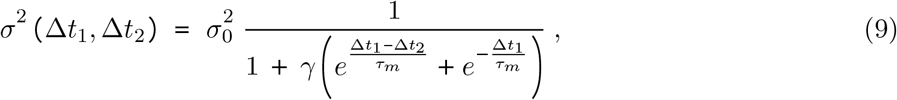

(see Supplementary Text A.2 for a derivation), where *σ*_0_ and *γ* are synaptic constants that scales the noise contribution to *u*.

The posterior PSC distribution *m* (Δ*t*_1_, Δ*t*_2_) and *v* (Δ*t*_1_, Δ*t*_2_) in Eq. (2) were computed for the LIF neuron model (Section 3.3), given by

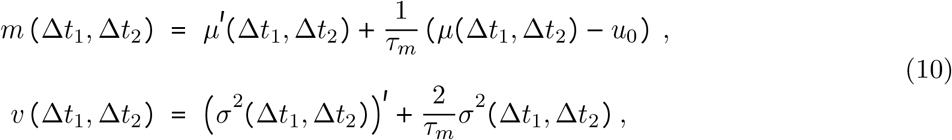

where 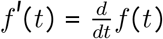 denotes the time derivative (see Supplementary Text A.3 for a detailed derivation).

### Numerical simulations

All simulations were performed in Python (3.8.5) using the Euler method to approximate the solution of the stochastic differential equations with a fixed time step of 1 ms. Post-Synaptic currents were created as described in (A.13) where Dirac delta pulses were approximated by 1 ms rectangular pulses and truncated at zero to avoid negative currents. If not stated otherwise we used a synaptic release parameter 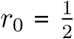. In Eq. (5) the membrane time constant *τ*_*m*_ was 30 ms, the resting potential *u*_0_ was -70 mV and the membrane resistance *R* was 10 MΩ. The firing threshold *ϑ* was -55 mV, *u*_*r*_ was -75 mV and the learning rate *η* was 10^−5^. In Eq. (9) we used 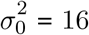 and *γ* = 50. Weights where drawn randomly and independently from a Gaussian distribution with mean and standard deviation of 10 and clipped at 0 before learning.

In Fig. 2 we used a single s-FEP synapses, applied different pre/post spike pairing protocols and recorded the resulting weight changes. In Fig. 2D STDP spike protocols were repeated 50 times with a time delay of 500 ms between two pairings. In Fig. 2E independent Poisson spike trains were presented to the synapses for 100 s. Fig. 2F the STDP protocol from Fig. 2D was repeated with different initial weights.

In Fig. 3 we used single neurons that received input from 300 afferent input neurons. Input neurons fired a dense syn-fire chain where every neuron was emitting a spike in exactly 1 ms during a 300 ms time window. The post-synaptic neuron was brought to fire at the end of this pattern. In Fig. 3F and G the output firing was determined by the intrinsic neuron dynamics after learning. In Fig. 3G output spike times were drawn from a Gaussian distribution with different standard deviations during learning.

In Fig. 4 spike patterns were generated by randomly drawing values from a beta distribution (*α* = 0.2, *β* = 0.8) for each input channel and multiplying these values with the maximum rate of 20 Hz. From these rate patterns individual Poisson spike trains were drawn for every pattern. During the learning phase the output neurons were clamped to fire 50 Hz Poisson spike trains during presentation of the preferred stimulus pattern and remain silent otherwise. Pattern presentations were interleaved with phases of 200 ms of zero spiking on all input channels. In the supervised scenario, for every output neuron one of the five patterns was selected as preferred stimulus. During training the activity of the output neurons was clamped to fire during the presentation of the preferred stimulus pattern.

In the unsupervised scenario, the network was augmented with fixed lateral inhibition to stabilize the firing behavior during learning. All excitatory neurons were connected to the inhibitory neuron with a synaptic efficacy of 1. The inhibitory unit projected back to all excitatory neurons with a synaptic efficacy of -5. During learning output neurons were allowed to run freely according to their intrinsic dynamics. The s-FEP learning rule was used at all synapses between input and output neurons in both scenarios. In Fig. 4E we set the synaptic release parameter *r*_0_ to 1 to disable synaptic noise (‘*without noise*’ condition).

In Fig. 5 we used a recurrent network with 400 feedback neurons and 400 internal state neurons from which we selected 200 action neurons. Preferred positions of the feedback neurons where scattered uniformly over the action space and firing rates were scaled by the euclidean distance between agent position and preferred position. Action neurons were randomly assigned to preferred movement directions. Internal state neurons received lateral inhibitory feedback and rate adaptation as in Fig. 4. During spontaneous movement the agent’s end effector was set to *x*_*start*_ at trial onset, and then the position was updated every 50 ms by adding the decoded position offset provided by the action neurons (light blue traces in Fig. 5). During training the activity of action neurons was clamped to a pre-defined training trajectory (dark blue in Fig. 5).

## Acknowledgements

Written under partial support by the European Commission, Horizon 2020 Framework Programme for Research and Innovation, Grant Agreement No. 899265 (ADOPD). The authors would like to thank Anand Subramoney for valuable comments to the manuscript.

## A Supplementary information

In this Supplementary Text we provide the details to the derivation and implementation of the s-FEP model. This document is organized as follows: In Supplementary Text A.1 we review the main idea behind the FEP and how it is utilized here on the level of single synapses. In Supplementary Text A.2 we define the *generative density* that is used by the synapse to estimate the state of the somatic membrane potential. In Supplementary Text A.3 we define the *recognition density* that determines the dynamics of the stochastic synapse model. In Supplementary Text A.4 we develop our main theoretical result to show that the synaptic efficacy updates (3) of the main text, minimize the free energy *ℱ* (*z, w*) of the synaptic efficacy *w* with respect to the back-propagating action potentials *z*. In Supplementary Text A.5 we show that the same learning rules also emerge if the s-FEP is applied to a learning scenario for recurrent neural networks with arbitrary numbers of neurons and synapses.

### A.1 Synapse-level free energy minimization

Here we provide a brief overview over the main aspects of the FEP that are needed for our treatment of single synapses. Since we focus here on a relatively simple physical system – individual synapses that interacts with their post-synaptic neuron – we only need a subset of the theoretical framework that is provided by the FEP. An excellent comprehensive review on this topic can be found in [Buckley et al., 2017].

The FEP is a generic theoretical framework to describe the interaction of a behaving agent with its environment. Its main assumption is that the agent and the environment have physically separated states, the *internal* (*w*) and *external* (*u*) states, respectively, that cannot directly influence another. Interaction only takes place through specific *actions* (*y*), performed by the agent and *feedback* (*z*) provided by the environment. The FEP suggests that the agent should adapt its behavior to minimize the surprise caused by the feedback *z*, measured by the negative log likelihood, surprise (*z*) = − log *p* (*z*).

The FEP proposes a specific method to approaching a state of minimum surprise. This method rests on the idea that a biological organism maintains an internal model of its environment, that allows it to reason about the external states *u*. The internal model is composed of two parts, (1) the *recognition density q* (*u* | *w*), that describes how the external state *u* interacts with the internal state *w*, and (2) the *generative density p(u, z)*, that describes the dependency between external states *u* and feedback *z* [Buckley et al., 2017]. To simplify the notation we employ here the commonly used shortcut *q*_*w*_ (*u*) for (*q u* | *w*). The recognition density is parameterized by the internal state *w*. The generative density depends here on the set of somatic parameters that comprise the firing threshold *ϑ*, reset- and resting potential, *u*_*r*_ and *u*_0_, and the membrane time constant *τ*_*m*_. We assume that these parameters are constant and encoded *a-priori* into the dynamics of the synapses such that the dynamics of the soma and the synapse match (e.g. through evolutionary processes or adaptation that is significantly slower than the learning dynamics). Plasticity mechanisms to fine-tune the synaptic behavior to track changes in somatic parameters could be derived from the s-FEP framework as well, but are not the focus of this study.

Using this internal model the complexity of the surprise minimization problem can be approached by replacing the goal to minimize surprise directly by a variational upper bound, that allows us to split the problem into two parts. The theory stems from the observation that an upper bound on the surprise can be reached indirectly by employing the recognition density *q* to *guess* external states *u*, and the generative density *p* evaluates how well the feedback *z* agrees with the guessed external states *u*. The problem to minimize surprise is then augmented with a divergence term to also minimizing the mismatch between *q* and *p*.

We adopted this idea and suggested to minimize an upper bound on the surprise in every synapse, given by the variational free energy *ℱ*, which is defined as

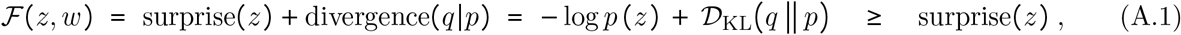

where *𝒟*_KL_(*q* ‖*p*) is the Kullback-Leibler (KL)-divergence between *q*_*w*_(*u*) and *p* (*u* | *z*). The inequality in (A.1) follows from *𝒟*_KL_(*q*‖*p*) ≥ 0 for any two probability distributions *q* and *p*. Inserting the definition of the KL-divergence we get

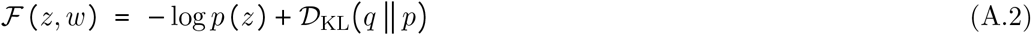

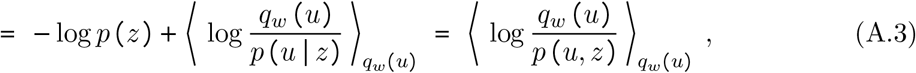

where 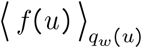 denotes the expectation of some function *f(u)* with respect to the probability density *q*_*w*_(*u*). By rearranging the terms of this last form, we can establish a link to the Helmholtz free energy that measures the useful energy potential in closed thermodynamic systems, from which the FEP inherits its name [Buckley et al., 2017,Neal and Hinton, 1998], by interpreting *ϵ* (*u, z*) = − log *p* (*u, z*) as the energy of state (*u, z*), to get

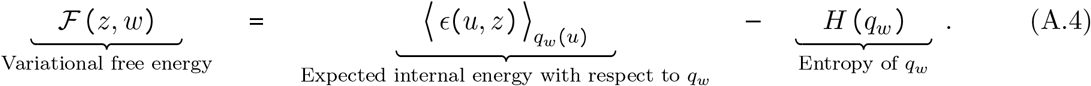

Using these definitions, we identify the relevant variables in our synapse model that are required by the FEP: (1.) *the internal states*, (2.) *the actions*, (3.) *the external states*, and (4.) the *feedback* (see [Friston, 2008] and Fig. 1 for an illustration).

1. *The internal states* summarizes all relevant internal variables that determine the behavior of the synapse. Since we focus here on long-term synaptic plasticity the internal state is given by the synaptic efficacy *w*. The internal states can be augmented with additional variables to also include other mechanisms, e.g. short term plasticity, but we neglect these here for the sake of simplicity.
2. *The actions* are utilized by the synapses to interact with the environment (the efferent neuron). In our model this is done through stochastic synaptic currents *y*, where the mean and variance of *y* is governed by the synaptic efficacy *w*. In our model, *y* denotes a sequence of synaptic currents (*y* = *y* (*t*) I *t* ≥ 0).
3. *The external states*. From the perspective of a synapse, the environment, it can immediately interact with, is the post-synaptic neuron. Here, we model the external states as the somatic membrane potential *u*(*t*) of a leaky integrate and fire (LIF) neuron with firing threshold *ϑ* and resting potential *u*_0_. We denote the whole sequence of the somatic membrane potential by (*u* = *u*(*t*) | *t* ≥ 0.
4. *The feedback*. In our model, a synapse only receives the back-propagating action potential of the post-synaptic neuron *z* as feedback to be informed about the somatic membrane potential. Formally, the spike train *z* is denoted by the set of firing times 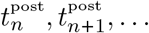 of the post-synaptic neuron. This feedback information about the external state *u*(*t*) is used by the synapse to update the internal model of the environment *p* (*u, z*).

Learning is realized in the FEP by minimizing *ℱ* (*z, w*) with respect to *w*, which can be done by gradient decent

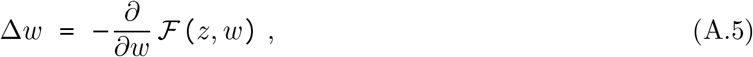

In the following sections we will derive the learning rule that solves this optimization problem for the case of our synapse model step by step. We consider the general form of the weight changes Δ*w* to show that the learning problem (A.5) can be solved by applying weight updates that only depend on the pre- and post-synaptic firing times, the current value of the synaptic efficacy and constants that are independent of learning, given by

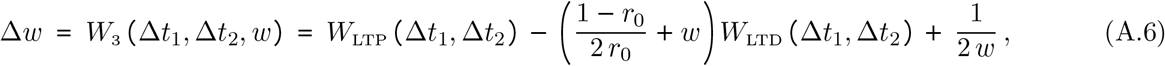

with 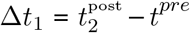 and 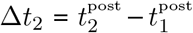. (A.6) is the general case for an arbitrary synaptic parameter *r*_0_. Eq. (3) shows the special case for 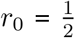 which was used throughout the paper (except Fig.4E). In our simulations we performed synaptic efficacy updates *w*_*new*_ = *w*_*old*_ + *η* Δ*w* for every post-pre-post spike triplet, with 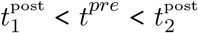, where 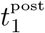 and 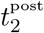 are the spike times of two neighboring post-synaptic spikes, and *t*^*pre*^ is a pre-synaptic spike time. *η* is a small positive constant learning rate *η* = 10^−5^. Weight updates were applied immediately at the arrival of the bAP events 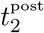 (no batching or buffering).

### A.2 The generative density

In this section we formally define the generative density *p* (*u, z*) which describes the joint dynamics of the membrane potential *u*, and the observed post-synaptic spike train *z* back-propagating to the synapse, in Eq. A.3. To arrive at this result it is convenient to think of the somatic membrane dynamics in terms of a deterministic function *g*, with *u*(*t*) = *g* (*y, t*), that maps a given sequence of synaptic input currents, *y*, to the current value of the membrane potential at time *t. g* is a piece-wise continuous function that obeys the membrane dynamics (5)-(7). The probability density of a membrane potential trace *u*, then reduces to a point density where *u* equals *g* at all times, i.e.

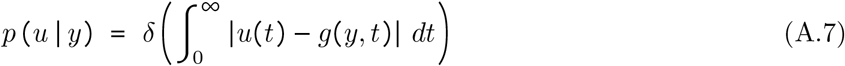

where *δ* is the Dirac delta function. We will further use the notation *u* = *g*(*y*) to denote the sequence of membrane potential values *u*(*t*) = *g*(*y, t*), for a given *y*.

Equation (A.7) can be used to assign a membrane potential trace *u* to a sequence of PSCs *y* with absolute certainty. However, a synapse does not have access to the true value of *y* (or *u*) since it may include input from other synapses that cannot be directly observed (and possibly further unobserved processes other than synaptic input). To reflect this uncertainty about *u* we assume for the definition of the generative density, that *y* is given by a stochastic process. Using this we can rewrite the dynamics of the membrane potential *u*(*t*), (5) in terms of a stochastic differential equation, by replacing the input current *y*(*t*), to get

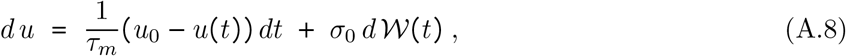

with resting membrane potential *u*_0_ and where *σ*_0_ scales the contribution of the total stochastic input and *d𝒲*(*t*) are the increments of the Wiener process.

(A.8) is an Ornstein-Uhlenbeck (OU)-process that describes the dynamics of the LIF neuron model with stochastic inputs [Gerstner et al., 2014]. This model is convenient because it compactly captures the uncertainty of a synapse that is not able to directly observe *u*(*t*) and all inputs to the post-synaptic neuron. The OU process can be solved analytically using stochastic calculus, e.g. if the process (A.8) is fixed to *u*_0_ at time 0 it evolves according to

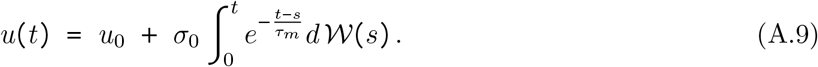

For long observation times the OU process converges to a stationary distribution, given by a Gaussian with mean *u*_0_ and variance 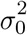, i.e. for 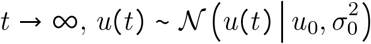.

The OU process (A.8) dynamics can be used to define the generative density *p* (*u, z*) for a LIF neuron. In the derivation of the s-FEP we make use of the fact, that also the posterior density *p* (*u* | *z*) can be solved explicitly. The information about the spike times *z* deflects the distribution of likely values of the membrane potential from its resting state, which is expressed in the posterior density *p* (*u* | *z*). We can express the dynamics of *u*(*t*) given the information that the membrane potential is at the firing threshold *ϑ* at the firing times *t*^post^, i.e. the constraint 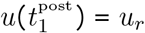 and 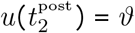 through a stochastic process with time varying mean *µ (t)* and variance *σ*^2^ (*t)*. Hence, we use a Gaussian process model of the external state, such that the posterior density is given by

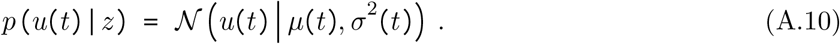

Using the FEP theory we can in principle assume any function *µ (t)* and *σ*^2^ (*t)* and develop learning rules that will strive to best approximate its dynamics. However, a reasonable choice will obey the constraints imposed by the neuron and synapse dynamics, e.g. the membrane time constant and the firing mechanism and resetting behavior of the neuron.

For LIF neuron model (A.8) the resulting *constraint stochastic process* has to fulfill the following requirements

1. The mean *µ*(*t*) obeys 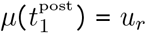 and 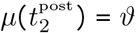.
2. For 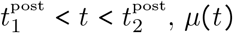 approaches the resting potential *u* asymptotically.
3. The variance *σ*^2^(*t*) obeys 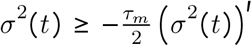 for 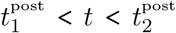, and approaches its minimum when close to the firing times 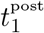 and 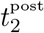.
4. For 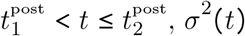 approaches the variance 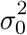 of the stationary distribution asymptotically.
5. The functions *µ(t)* and *σ*^2^ (*t)* are smooth and follow the LIF dynamics with time constant *τ*_*m*_. Constraint 3. incorporates that in the LIF dynamics, the variance can only shrink at a maximum speed proportional to the membrane time constant 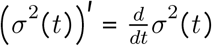 denotes the time derivative.

The LIF neuron implies OU process dynamics of the membrane potential. Given the information that the membrane potential is at the firing threshold *ϑ* at the firing times *t*^post^, i.e. the constraint 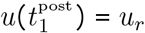 and 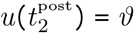, the OU process can be solved explicitly. The solution to this double constraint stochastic process is the OU-bridge process [Corlay, 2013, Szavits-Nossan and Evans, 2015]. For any neighboring postsynaptic spike pair 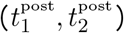 and time point *t* with 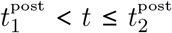 we can describe the dynamics of *u*(*t*) using its mean *µ*(*t*) and variance *σ*^2^(*t*). Using this result, for any neighboring postsynaptic spike pair 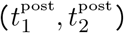 and time point *t* with 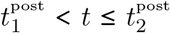 we describe the dynamics of *u*(*t*) using the mean *µ*(*t*) and variance function *σ*^2^(*t*), given by

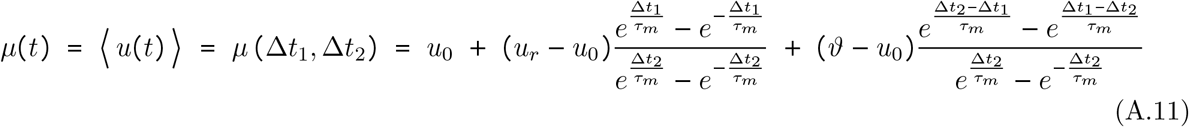

and

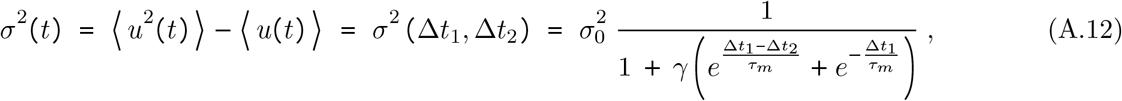

where 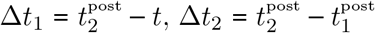 and *γ* is a constant that scales the slope of the variance function. In other words, the dynamics of the membrane potential subject to the constraint 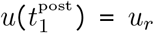 and 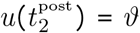 are described by a stochastic process with mean *µ (t)* and variance *σ*^2^ (*t)*. The membrane potential mean and variance functions (A.11) and (A.12) are piece-wise defined for all postsynaptic spike intervals 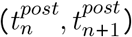. The membrane dynamics during each interval are statistically independent of each other due to the resetting behavior of the neuron model. In all simulations we used *γ* = 50 and 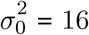.

The mean function *µ (t)* in (A.11) is identical to the OU-bridge process model [Corlay, 2013, Szavits-Nossan and Evans, 2015]. This function describes the asymptotic approach to the resting potential *u*_0_ and the slope towards the firing threshold *ϑ*. The variance function *σ*^2^ (*t)* in (A.12) is flatter than the direct solution of the OU-bridge process model to incorporate the additional constraint 3.

Using the definition (A.11) and (A.12), we find that the posterior of the generative density can be evaluated at any time point *t*, as *p* (*u*(*t*) | *z*) = *𝒩* (*u*(*t*) | *µ*(*t*), *σ*^2^(*t*)). Furthermore, since the mean (A.11) and variance (A.12) functions only depend on the relative spike timing Δ*t*_1_ and Δ*t*_2_ we find that these quantities can be expressed in the form *µ* (Δ*t*_1_, Δ*t*_2_) and *σ*^2^ (Δ*t*_1_, Δ*t*_2_).

### A.3 The recognition density

Here we define the recognition density *q*_*w*_ *u* for our synapse model. A synapse injects brief stochastic current pulses of amplitude proportional to the current value of the synaptic efficacy into the post-sypantic neuron when a pre-synaptic input arrives. For the derivation that follows we will use here the simplifying assumption that *y(t)* is given by the PSCs of a single synapse to keep the notation uncluttered. In Supplementary Text A.5 we will show that the same learning model also arises when the s-FEP is applied to a network of neurons with an arbitrary number of synapses.

Let the spike train of the pre-synaptic neuron be denoted by *z*_pre_(*t*), given by Dirac delta pulses centred at the pre-synaptic firing times. The post-synaptic input current *y*(*t*) is then given by

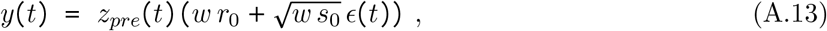

where *w* ≥ 0 is the synaptic efficacy, *z*_*pre*_ is a spike train given by Dirac delta pulses centered at pre-synaptic spike times, and *ϵ*(*t*) is a source of independent unit variance zero mean Gaussian noise. The constant *r*_0_ and *s*_0_ scale the mean and variance of the synaptic current. We used 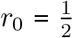 and *s*_0_ = *r*_0_ (1−*r*_0_) if not stated otherwise in accordance with previous models [Katz, 1971].

The stochastic synapse model (A.13) suggests that at the time points of pre-synaptic firing *t* = *t*^*pre*^ the amplitudes of synaptic currents follow a Gaussian distribution. To make this explicit we rewrite (A.13) to get

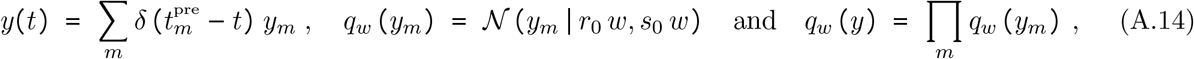

where the sum runs over all pre-synaptic firing times. This Gaussian model, (A.14), is an approximation to previous models of stochastic synaptic release [Katz, 1971].

At this point it is instructive to note that the generative density, that was introduced in Supplementary Text A.2, can be solved directly for *y*. More precisely, we will show that *p(y|z)* can be expressed as a Gaussian distribution with time-varying mean and variance functions *m*(*t*) and *v*(*t*), such that *y*_*m*_ ∼*𝒩* (*y*_*m*_ | *m*(*t*_*m*_), *v*(*t*_*m*_)). Therefore, the generative density can be inverted, yielding a time varying distribution over PSCs *y*(*t*) that, when injected into the post-synaptic neuron, will give the desired distribution for the membrane potential dynamics. This result can be obtained by stochastic integration, but to keep this paper self-contained we provide a proof here. We start by considering a general drift-diffusion process and then show the special case of the LIF neuron dynamics step by step.

In general, the evolution of the probability density function *p*(*u, t*) of a stochastic process *u* at time *t*, with drift *A*(*u, t*) and diffusion *B*(*u, t*), can be described by the Fokker-Planck equation

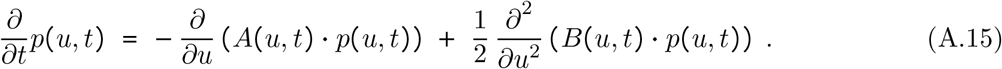

Note that *u* denotes here an instantaneous value rather than whole sequences. To describe the dynamics of our model neuron we treat here the case that *p*(*u, t*) is a Gaussian distribution with time-varying mean *µ*(*t*) and variance *σ*^2^(*t*) functions, i.e. *u*(*t*) ∼ *𝒩* (*u*(*t*)|*µ*(*t*), *σ*^2^(*t*)) at any time point *t*, to get for the left side of (A.15)

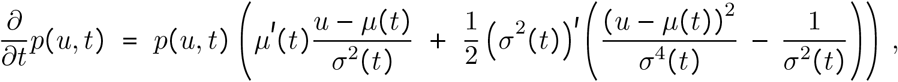

where 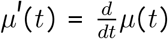 and 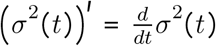 denote the time derivatives. Furthermore we can expand the right side of (A.15) to get

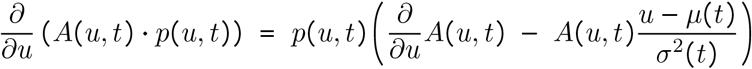

and

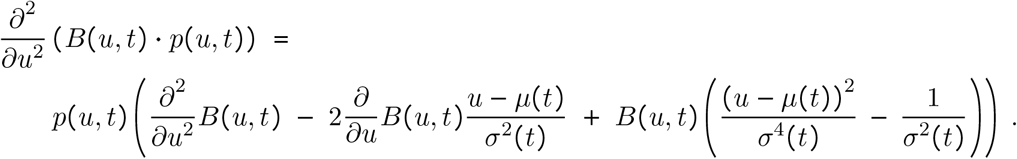

Therefore, by plugging these results back into the Fokker-Planck equation (A.15), we find the condition that has to be satisfied for functions *A (u, t)* and *B (u, t)*to be given by

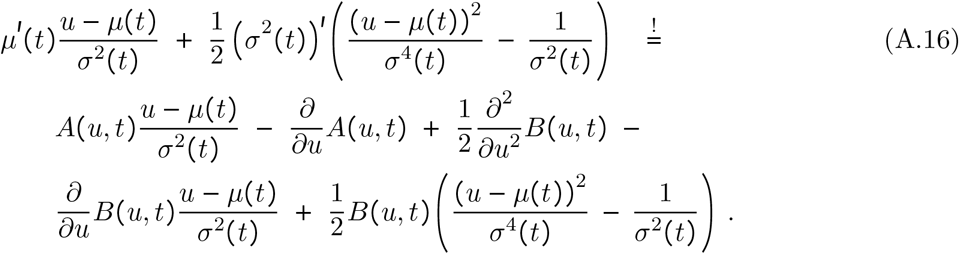

Clearly, the choice *A*(*u, t*) = *µ*^′^(*t*), *B*(*u, t*) = (*σ*^2^(*t*))^′^ satisfies this condition for any differentiable functions *µ (t)* and *σ*^2^ (*t)*.

This last results holds in general. To arrive at the final result we replace the general drift-diffusion dynamics with the special case of a leaky integrator with finite integration time constant *τ*_*m*_ using the ansatz 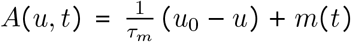 and *B (u, t) = v (t)*. In this case, we can make condition (A.16) satisfied if 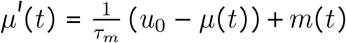 and 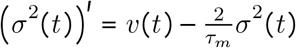. This can be verified by plugging this result back into (A.16)

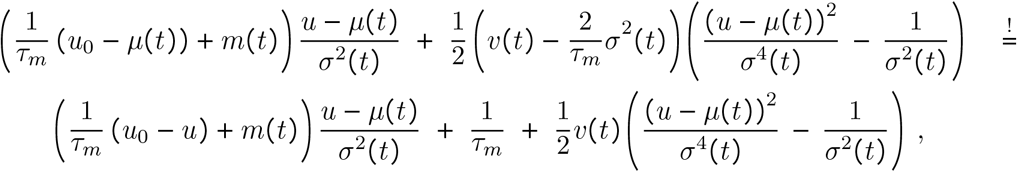

from which the equality follows

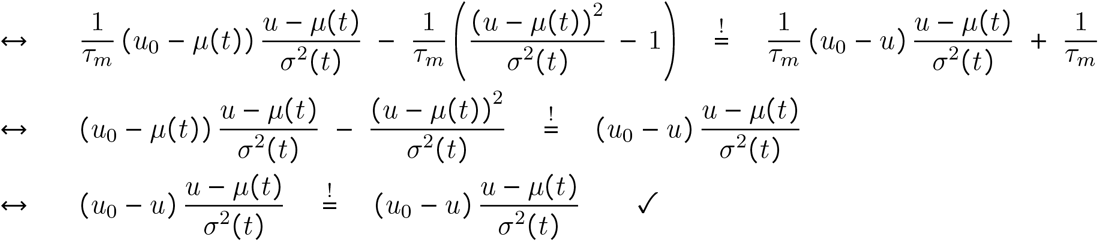

This proofs that a stochastic process *u* with 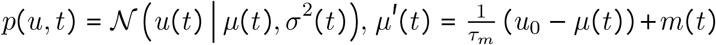 and 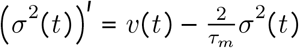 is realized by a drift 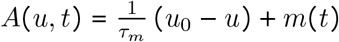 and diffusion *B*(*u, t*) = *v(t)*. Equivalently, any process *u* with mean *µ (t)* and variance *σ*^2^(*t*) can be realized if

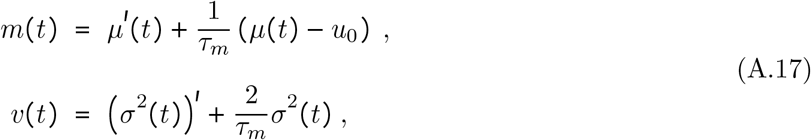

and *v (t)* ≥ 0 can be satisfied for all *t*. This last result is used in Supplementary Text A.4 to derive the learning rule (A.6).

Furthermore, by integration of this last result we find that any integrable functions *m (t)* and *v (t)* > 0 yield the following dynamics for the stochastic process *u*

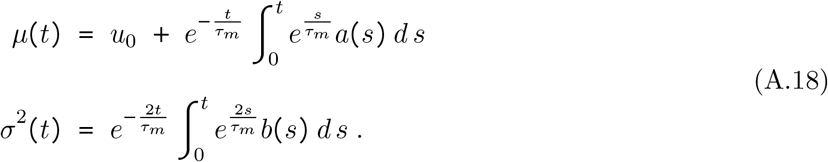

For *m*(*t*) = 0 and *v*(*t*) = *b* (constant) we recover the Ornstein-Uhlenbeck process dynamics.

Therefore, we define the posterior density *p* (*y* | *z*) as the distribution over *y* where any instantaneous value *y*_*m*_ at time *tm obeys*

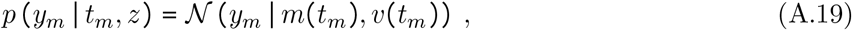

where *m*(*t*) and *v*(*t*) are as defined in (A.17) with *µ*(*t*) and *σ*^2^ (*t*) given by the solution to *p* (*u* | *z*) as defined in (A.11) and (A.12), respectively. Clearly we can recover *p* (*u* | *z*) using (A.7) by marginalizing over *y*

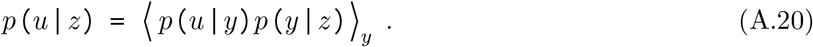

### A.4 Derivation of the learning rule

Next we make use of the result from Supplementary Text A.2 and Supplementary Text A.3 to develop the learning rules that minimize the variational free energy (A.3). The approach that is taken here is very similar to [Kingma and Welling, 2013] and makes extensive use of the fact that the generative density *p (u, z)* can be expressed analytically and Bayesian posteriors can be solved in closed form. We also assume that the neuron parameters, that determine the generative density, are constant and known *a priori*, and hence do not need to be learned by the synapses (however learning rules to infer these parameters could be easily derived from the model, but are not the focus of this study). Also, other than previous applications of the FEP (e.g. [Isomura et al., 2016]), we assume that the generative density *p (u, z)* is independent of *w*. While this assumption is mathematically sound and commonly made in applications of variational Bayesian methods (e.g. [Mnih and Gregor, 2014]) it is unusual in the FEP literature. In Supplementary Text A.5 we provide an additional justification of this choice for the s-FEP.

Using the assumptions and definitions outlined above, (A.5) becomes

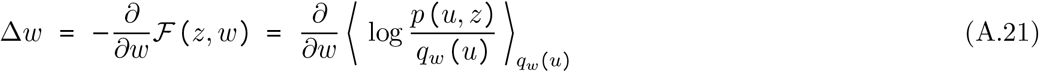

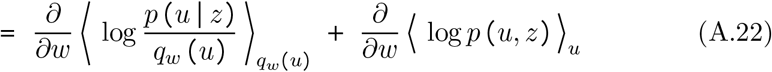

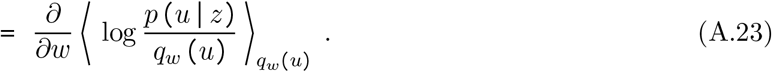

The first equality follows from Bayes rule and the second follows since ⟨ log *p (u, z)* ⟩ _*u*_ is constant in *w* by construction. Next, we exploit here that the OU process model (A.8) suggests a one-to-one relation between PSC traces *y* and somatic membrane potentials *u*, that is, for a given *y* we can determine *u* through a deterministic function. For the deterministic function *u* = *g(y)*, we can replace the marginal over *q*_*w*_ (*u*) in (A.23) with a marginal over *q*_*w*_ (*y*), i.e.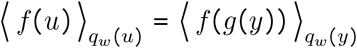. This is a variant of the reparameterization trick [Kingma and Welling, 2013] that reflects here the fact that at the post-synaptic spike times *t*^post^, *p (u, z)* collapses to a point mass density.

Using this result we get

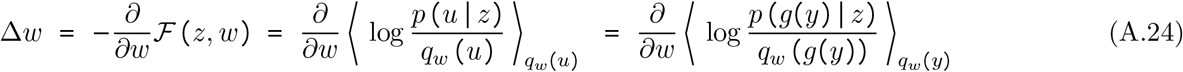

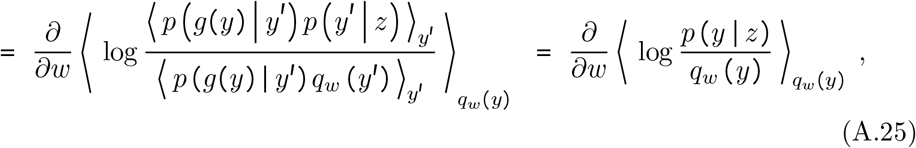

where in the last equality we used Eq. A.7. Thus, the minimization of the variational free energy (A.1) with respect to the synaptic efficacy *w* of a synapse that interacts with its post-synaptic neuron, can be reduced to a gradient on the mismatch between the posterior synaptic current *y* given *z* and the recognition density *q*_*w*_(*y*). Finally, we use that the PSC pulses in *y* are independent and thus the marginal in (A.25) can be replaced by a sum over pre-synaptic firing times 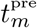

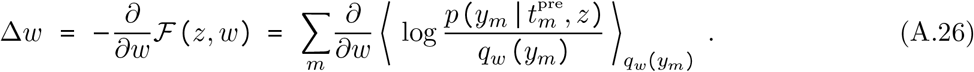

Also note, that it is sufficient here to consider only the two post-synaptic spikes in *z*, that are directly neighboring 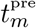, to evaluate 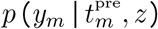 since the somatic membrane dynamics are independent after the reset to *u*_*r*_.

This result is useful, because the generative model established in Supplementary Text A.2 allows us to express the posterior distribution *p*(*y*_*m*_ | *t*_*m*_, *z*) in closed form. In Supplementary Text A.3 we show in detail that a synaptic current *y* ∼ *𝒩* (*y* | *m*(*t*_*m*_), *v*(*t*_*m*_)) enables a synapse to realize a somatic membrane potential *u*(*t*) that obeys the stochastic processes with mean *µ*(*t*) and variance *σ*^2^(*t*), if

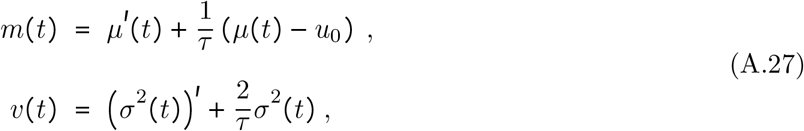

where *µ*(*t*) and *σ*^2^(*t*) are as defined in (A.11) and (A.12).

To construct the term 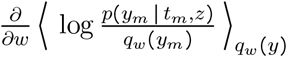 of (A.26) we use the result from Supplementary Text A.2 and assume a general Gaussian form 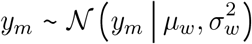 to get

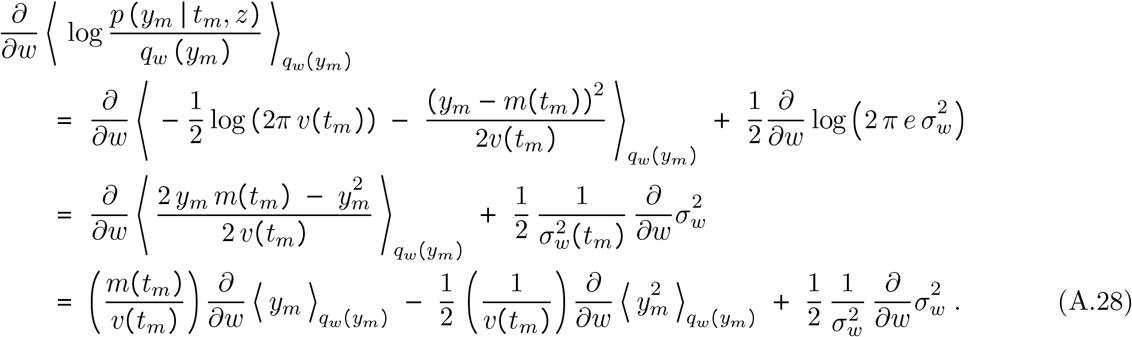

By plugging in (A.11) and (A.12) we recover the LTP and LTD term in (A.6).

Using 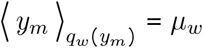 and 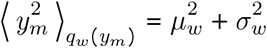, we get

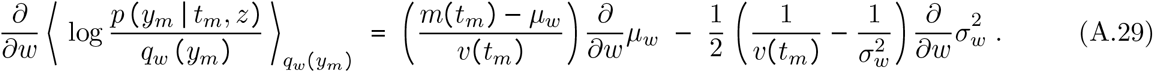

Finally, using (A.13) we identify *µ*_*w*_ and 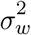 to get the result for any *t*_*m*_ at the pre-synaptic firing times

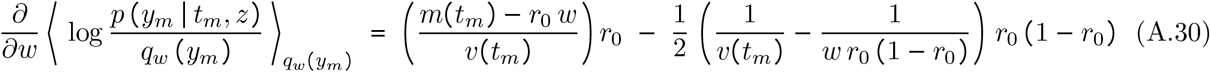

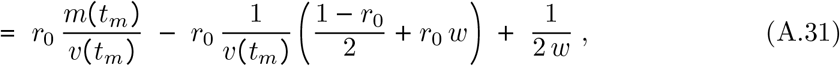

which is identical to the result in (A.6) with 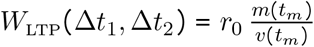 and 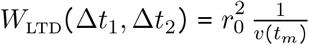.

Using this we identify the triplet STDP windows, given by

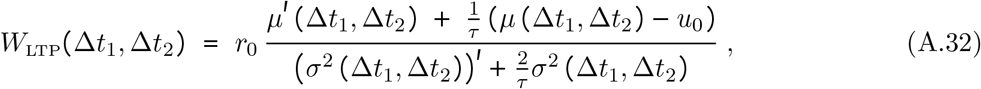

and

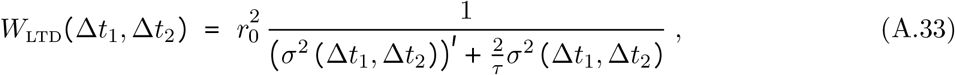

where *µ* (Δ*t*_1_, Δ*t*_2_) and *σ*^2^ (Δ*t*_1_, Δ*t*_2_), respectively, are the estimated mean and variance of the membrane potential based on the back-propagating action potentials ((A.11) and (A.12)), and 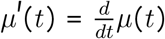 and 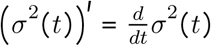 denote the time derivatives.

The rational underlying the learning model is illustrated in Fig. 2A. For any pre-synaptic firing time *t*_*m*_ a random PSC is generated using the recognition density *q*_*w*_ (*y*_*m*_). In addition post-pre-post spike triplets are formed by considering the neighbouring post-synaptic spikes back-propagating to the synapse. Based on these spike triplets, the posterior density *p(y*_*m*_|*t*_*m*_, *z*) is constructed and the mismatch with the recognition density triggers a weight update. The internal model does not need to be represented explicitly but is implicit in the shape of the triplet STDP learning window.

The proposed s-FEP learning rules installs a single parameter distribution *q*_w_ (*y*) that minimizes the distance to the two-parameter posterior density *y* ∼ *𝒩* (*y* | *m*(*t*), *v*(*t*)). In Fig. 3E we showed that the synaptic efficacies are correlated with the euclidean norm of the posterior *µ*^∗^ and *σ*^∗^. Using the result (A.31) we can make a more careful analysis, by keeping the PSC posterior fixed, *m*(*t*) = *µ*^∗^ and *v*(*t*) = (*σ*^∗^)^2^, and then analyzing the roots of the learning rule (A.31). Using this we find that the weights converge to 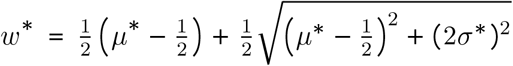, (with 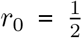as in our simulations).

Hence, weights encode both the target mean and variances, i.e. for small *σ*^∗^ and large *µ*^∗^ we have *w*^∗^ ≈ *µ*^∗^ and for large *σ*^∗^ and small *µ*^∗^ we have *w*^∗^ ≈ *σ*^∗^.

### A.5 Network level FEP emerges from the s-FEP

In the previous sections we have derived the s-FEP for a single synapse for the sake of illustration. Clearly this is a very limiting case and in general we are interested in a FEP that concerns arbitrarily large recurrent neural networks. In this section we thus turn to a network-level treatment of the s-FEP. We show that the same learning rules (A.23) also applies to a class of learning problems and networks of neurons with an arbitrary number of neurons and synapses. To show this we use here vectorized forms **z** = (*z*_1_, …, *z*_*K*_), **u** = (*u*_1_, …, *u*_*K*_), **y** = (*y*_1_, …, *y*_*K*_) and **w** = (*w*_11_, …, *w*_*KK*_) to denote ordered sets of network spikes, membrane potentials, PSCs and the synaptic efficacies, respectively, for networks of *K* neurons. *w*_*ki*_ denotes the synaptic efficacy from neuron *i* to neuron *k*.

In addition we assume that there is an additional external feedback **x** that may influence the activity of an arbitrary subset of network neurons. This feedback is assumed to be a sensory experience of some form that is directly perceptible by a subset of the neurons. The feedback may take the role of a teacher signal as in our example in Fig. 5, but it can also be a more abstract signal that indicates e.g. goal reaching or aversive stimuli. Subsequently we will take the free energy minimization problem to the network level by considering the network spikes **z** as part of the state space and minimizing the augmented variational free energy

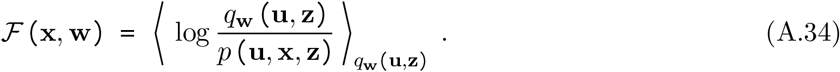

#### The network-level generative density

The corresponding generative density *p* (**u, x, z**) now captures the dependence between states (**u, z**) and feedback **x. x** can in principle be an arbitrary vectorvalued function of time, but as in the single synapse case we focus here on models where the posterior density can be solved and results in conditional independence between individual membrane potentials *u*_*k*_

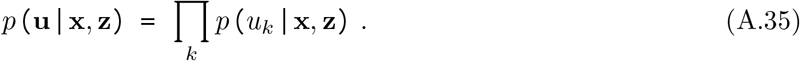

The following derivations will be valid for scenarios where this posterior density can be evaluated. **x** can for example be given by strong external inputs that impose a certain firing pattern on a subset of network neurons or a bias potential that offsets the resting potential of some neurons. Scenarios where the independence (A.35) does not hold lead to more complex learning rules that are not further studies here (see the discussion at the end of this section).

As in the single synapse case we can make use of another conditional independence that is imposed by the neuron’s resetting behavior at post-synaptic firing times. This implies conditional independence between any two post-synaptic spike times 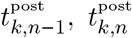. To make this explicit we denote by 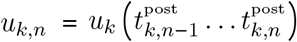 and 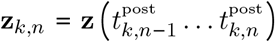 subsets of the sequences *u*_*k*_ and **z**, respectively, over the time interval 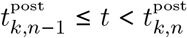. Using this we can write for (A.35)

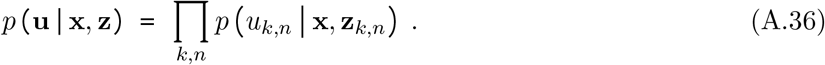

#### The network-level recognition density

For the single synapse model we assumed the pre-synaptic spikes to be given, which considerably simplified the derivations. This is not possible anymore for the network-level model, which also has to reflect the recurrent input from network spikes **z** from other neurons. To account for this additional dynamics we consider the network-level PSCs, *y*_*k*_(*t*) of neuron *k*

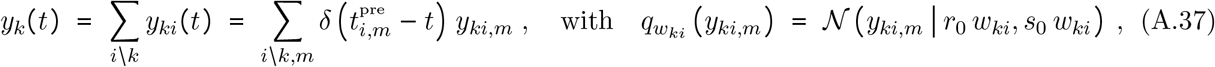

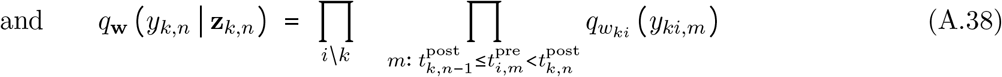

where the sum over *i* runs over all network neurons excluding *k* (fully recurrent network without autapses). *y*_*ki,m*_ denotes the PSC amplitude of the *m*-th PSC event under synapse *ki*, and *y*_*k,n*_ denotes the whole PSC sequence of neuron *k* in the time interval 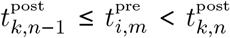. The pre-synaptic firing times 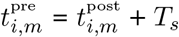, denote the recurrent network activity with some small time delay *T*_*s*_. Eq. (A.38) includes the implicit assumption that no pre-synaptic spikes arrive at neuron *k* at the exact same time 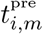 such that the conditional independence holds (further discussed at the end of this section).

The network-level recognition density determines the response of the network to the PSCs (A.38). To define the network-level recognition density, we can make use of the fact that the neuron model is Markovian, which allows us to define *q*_**w**_(**u, z**) in terms of a distribution over individual network spike times 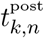

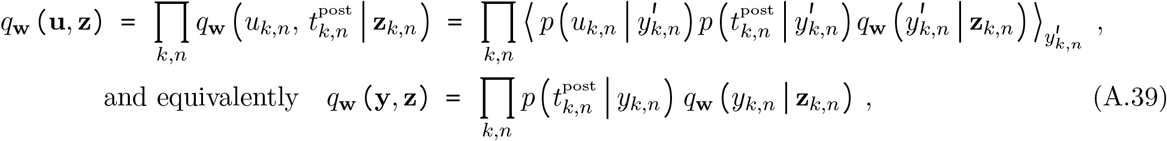

Where 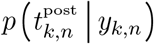 is the probability density over firing time 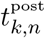 in response to given PSCs *y*_*k,n*_, given by a point density similar to (A.7), where the membrane potential reaches the firing threshold exactly at time 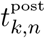.

Using this result we get for the network level synaptic efficacy updates

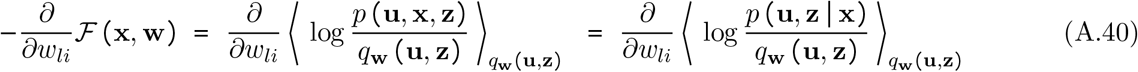

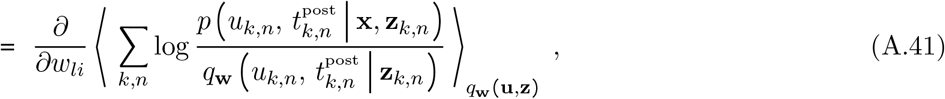

where in (A.40) we used that the derivative of ⟨ *p (***u, x, z)** ⟩ _**u**,**z**_ with respect to *w*_*li*_ is zero by construction. Again by exploiting the properties of (A.39) we can use reparameterization to replace the marginal over *u* with one over *y*. For (A.41) we get

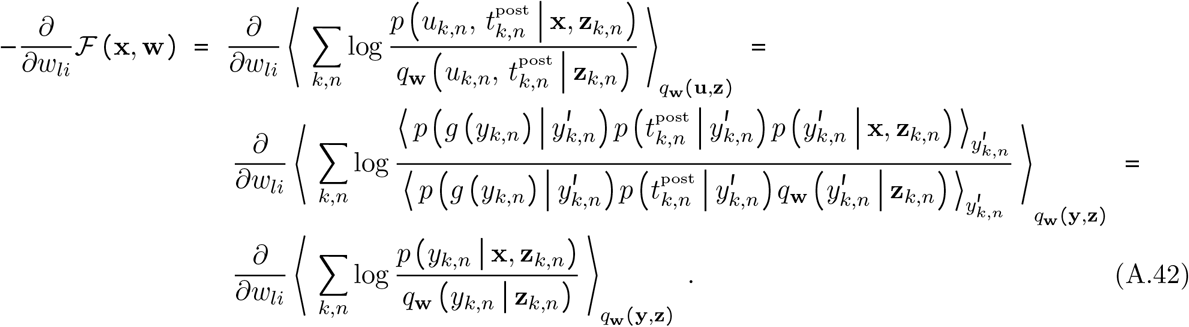

Note that the information about **z** is redundantly encoded in **u** and therefore this dependence vanishes under the expectation. Finally, using this result and (A.38), we identify the synaptic efficacy updates for *w*_*li*_

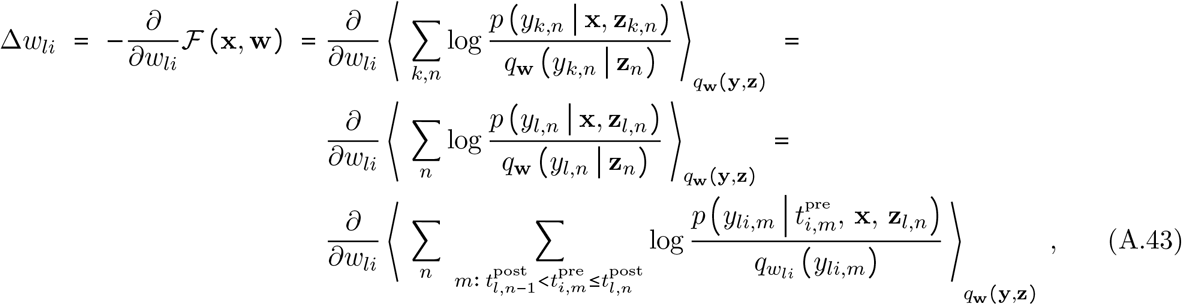

where 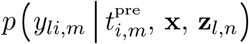 is the posterior density over PSCs evaluated at time 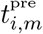.

A sampling-based approximation of (A.44) can be realized by first, sampling **y, z** ∼ *q*_**w**_(**y, z**) by letting the network evolve according to its intrinsic dynamics. Then, update the synaptic efficacies for every post-pre-post spike according to

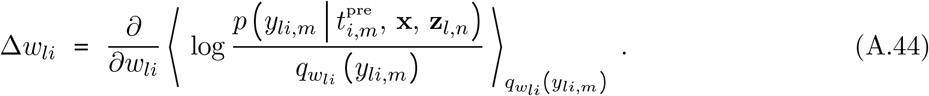

In summary we find that network-level learning can be established in the from (A.44) if the following assumptions hold

- Network spikes never arrive simultaneously so that the conditional independence in (A.38) applies.
- The posterior of the generative density 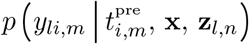 can be computed in the synapses (realized through the learning rules).

The first assumption is easy to fulfill for a general recurrent network since the theoretical framework leaves enough room to de-correlate input spikes. Note, that the probability that any two neurons spike exactly at the same time already approaches zero in continuous time if spikes are generated independently in every neuron. More complex cases like multiple input synapses from the same neuron can be treated by including a small independent jitter on the synaptic delays. In our simulations we found that simultaneous arrival of input spikes does not seem to be problematic in practice and no additional measures where taken to prevent them.

The second assumption is clearly only true for special cases where the implied conditional independence holds. In Fig. 4 and 5 we chose the most basic example for this to be true. There, we used a simple model where **x** provided external input to drive a subset of neurons to fire at externally defined time points (e.g. through external sensory inputs that provide strong inputs to some neurons). In this simple example 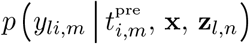 in (A.44) can be expressed analogously to Eq. (A.19) and the resulting learning rules are identical to (3) with the only difference that spike times are generated by the intrinsic dynamics *q*_**w**_(**y, z**) for neurons that are not driven externally by **x**. In more complex scenarios the posterior can be realized through additional learning mechanism, e.g. through neuromodulatory signals, as previously studied in [Isomura et al., 2016].

#### Reasons against a weight-dependent generative density for the s-FEP

Our derivations rely on the assumption that the generative density is independent of the synaptic efficacies **w**. This implies that the generative- and the recognition density have separate parameter spaces. While this assumption is mathematically sound and has been used in many previous applications of variational Bayes (e.g. [Mnih and Gregor, 2014]) it is less common in the FEP literature. However, we argue here that in the context of the s-FEP, a weight-dependent generative density *p*_**w**_ **(u, x, z)** would not be beneficial computationally and also would not improve biological realism.

On the computational side, we find the main difference that the term 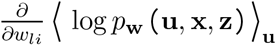 would not vanish in (A.41) and therefore this gradient has to be estimated in every synapse. It is well known that this kind of gradients is notoriously hard to compute and suffers from adverse signal-to-noise properties [Mnih and Gregor, 2014]. For the case of the s-FEP the situation is even worse because the learning signal has to be estimated based on the sparse bAPs that arrive at the synapse. The effect of a single synapse onto the somatic membrane potential is so minuscule that detecting this effect using a sampling-based estimator would require an unreasonable number of samples before it converges to a meaningful estimate of the gradient. An explicit dependence between the s-FEP generative density of the somatic membrane potential and all synaptic efficacies does not seem to be biologically feasible either, because it would require every synapse in a neuron to have knowledge about the PSCs of all other synapses.

For these reasons we decided to pursue a model-based, bottom-up approach to estimate the s-FEP gradient by splitting the parameter space of the recognition- and the generative density. Therefore, synapses maintain only a minimal model of the soma that reflects the high uncertainty of the synapse due to the sparse feedback. This has the advantage that the solution of the s-FEP can be expressed directly in closed form and manifests in learning rules that only require variables that are locally available at the synapse. Our derivation does not require further approximations such as linearizations to develop this result. Despite the simplicity of the generative model that is locally maintained in every synapse, we show in simulations (Fig. 4 and 5) and theory (Supplementary Text A.5) that the s-FEP can be scaled up to the level of recurrent networks.

## References

Aitchison, L., Jegminat, J., Menendez, J. A., Pfister, J.-P., Pouget, A., and Latham, P. E. (2021). Synaptic plasticity as bayesian inference. Nature Neuroscience, 24(4):565–571.

Aitchison, L., Pouget, A., and Latham, P. E. (2014). Probabilistic synapses. arXiv preprint 1410.1029.

Barascud, N., Pearce, M. T., Griffiths, T. D., Friston, K. J., and Chait, M. (2016). Brain responses in humans reveal ideal observer-like sensitivity to complex acoustic patterns. Proceedings of the National Academy of Sciences, 113(5):E616–E625.

Bastos, A. M., Usrey, W. M., Adams, R. A., Mangun, G. R., Fries, P., and Friston, K. J. (2012). Canonical microcircuits for predictive coding. Neuron, 76(4):695–711.

Borst, J. G. G. (2010). The low synaptic release probability in vivo. Trends in neurosciences, 33(6):259–266.

Brea, J., Senn, W., and Pfister, J.-P. (2013). Matching recall and storage in sequence learning with spiking neural networks. The Journal of Neuroscience, 33(23):9565–9575.

Buckley, C. L., Kim, C. S., McGregor, S., and Seth, A. K. (2017). The free energy principle for action and perception: A mathematical review. Journal of Mathematical Psychology, 81:55–79.

Buesing, L. and Maass, W. (2008). Simplified rules and theoretical analysis for information bottleneck optimization and pca with spiking neurons. In Advances in Neural Information Processing Systems, pages 193–200.

Buesing, L. and Maass, W. (2010). A spiking neuron as information bottleneck. Neural computation, 22(8):1961–1992.

Caporale, N. and Dan, Y. (2008). Spike timing–dependent plasticity: a hebbian learning rule. Annu. Rev. Neurosci., 31:25–46.

Chalk, M., Marre, O., and Tkačik, G. (2018). Toward a unified theory of efficient, predictive, and sparse coding. Proceedings of the National Academy of Sciences, 115(1):186–191.

Corlay, S. (2013). Properties of the ornstein-uhlenbeck bridge. arXiv preprint 1310.5617.

Cornejo, V. H., Ofer, N., and Yuste, R. (2021). Voltage compartmentalization in dendritic spines in vivo. Science, page eabg0501.

Dan, Y. and Poo, M.-m. (2004). Spike timing-dependent plasticity of neural circuits. Neuron, 44(1):23–30.

Davies, M., Srinivasa, N., Lin, T.-H., Chinya, G., Cao, Y., Choday, S. H., Dimou, G., Joshi, P., Imam, N., Jain, S., et al. (2018). Loihi: A neuromorphic manycore processor with on-chip learning. Ieee Micro, 38(1):82–99.

Deneve, S. (2008). Bayesian spiking neurons ii: learning. Neural computation, 20(1):118–145.

Driscoll, L. N., Pettit, N. L., Minderer, M., Chettih, S. N., and Harvey, C. D. (2017). Dynamic reorganization of neuronal activity patterns in parietal cortex. Cell, 170(5):986–999.

Friston, K. (2005). A theory of cortical responses. Philosophical transactions of the Royal Society B: Biological sciences, 360(1456):815–836.

Friston, K. (2008). Variational filtering. NeuroImage, 41(3):747–766.

Friston, K. (2010). The free-energy principle: a unified brain theory? Nature reviews neuroscience, 11(2):127.

Friston, K. (2012). A free energy principle for biological systems. Entropy, 14(11):2100–2121.

Friston, K., Schwartenbeck, P., FitzGerald, T., Moutoussis, M., Behrens, T., and Dolan, R. J. (2014). The anatomy of choice: dopamine and decision-making. Philosophical Transactions of the Royal Society B: Biological Sciences, 369(1655):20130481.

Friston, K., Thornton, C., and Clark, A. (2012). Free-energy minimization and the dark-room problem. Frontiers in psychology, 3:130.

Gerstner, W., Kistler, W. M., Naud, R., and Paninski, L. (2014). Neuronal dynamics: From single neurons to networks and models of cognition. Cambridge University Press.

Gjorgjieva, J., Clopath, C., Audet, J., and Pfister, J.-P. (2011). A triplet spike-timing–dependent plasticity model generalizes the bienenstock–cooper–munro rule to higher-order spatiotemporal correlations. Proceedings of the National Academy of Sciences, 108(48):19383–19388.

Gontier, C. and Pfister, J.-P. (2020). Identifiability of a binomial synapse. Frontiers in computational neuroscience, 14:86.

Graupner, M. and Brunel, N. (2012). Calcium-based plasticity model explains sensitivity of synaptic changes to spike pattern, rate, and dendritic location. Proceedings of the National Academy of Sciences, 109(10):3991–3996.

Grollier, J., Querlioz, D., Camsari, K., Everschor-Sitte, K., Fukami, S., and Stiles, M. D. (2020). Neuromorphic spintronics. Nature electronics, 3(7):360–370.

Harris, J. J., Jolivet, R., and Attwell, D. (2012). Synaptic energy use and supply. Neuron, 75(5):762–777.

Indiveri, G., Linares-Barranco, B., Legenstein, R., Deligeorgis, G., and Prodromakis, T. (2013). Integration of nanoscale memristor synapses in neuromorphic computing architectures. Nanotechnology, 24(38):384010.

Isomura, T. and Friston, K. (2018). In vitro neural networks minimise variational free energy. Scientific reports, 8(1):16926.

Isomura, T. and Friston, K. (2020). Reverse-engineering neural networks to characterize their cost functions. Neural Computation, 32(11):2085–2121.

Isomura, T., Kotani, K., and Jimbo, Y. (2015). Cultured cortical neurons can perform blind source separation according to the free-energy principle. PLoS computational biology, 11(12):e1004643.

Isomura, T., Sakai, K., Kotani, K., and Jimbo, Y. (2016). Linking neuromodulated spike-timing dependent plasticity with the free-energy principle. Neural computation, 28(9):1859–1888.

Jensen, T. P., Zheng, K., Cole, N., Marvin, J. S., Looger, L. L., and Rusakov, D. A. (2019). Multiplex imaging relates quantal glutamate release to presynaptic ca 2+ homeostasis at multiple synapses in situ. Nature communications, 10(1):1–14.

Jimenez Rezende, D., Wierstra, D., and Gerstner, W. (2011). Variational learning for recurrent spiking networks. In Neural Information Processing Systems.

Kanai, R., Komura, Y., Shipp, S., and Friston, K. (2015). Cerebral hierarchies: predictive processing, precision and the pulvinar. Philosophical Transactions of the Royal Society B: Biological Sciences, 370(1668):20140169.

Katz, B. (1971). Quantal mechanism of neural transmitter release. Science, 173(3992):123–126.

Kiebel, S. J. and Friston, K. J. (2011). Free energy and dendritic selforganization. Frontiers in systems neuroscience, 5:80.

Kingma, D. P. and Welling, M. (2013). Auto-encoding variational bayes. arXiv preprint 1312.6114.

Kostadinov, D., Beau, M., Pozo, M. B., and Häusser, M. (2019). Predictive and reactive reward signals conveyed by climbing fiber inputs to cerebellar purkinje cells. Nature Neuroscience, 22(6):950–962.

Levy, W. B. and Baxter, R. A. (1996). Energy efficient neural codes. Neural computation, 8(3):531–543.

Levy, W. B. and Baxter, R. A. (2002). Energy-efficient neuronal computation via quantal synaptic failures. Journal of Neuroscience, 22(11):4746–4755.

Linsker, R. (1988). Self-organization in a perceptual network. Computer, 21(3):105–117.

Maass, W. (2014). Noise as a resource for computation and learning in networks of spiking neurons. Proceedings of the IEEE, 102(5):860–880.

Mayr, C., Hoeppner, S., and Furber, S. (2019). Spinnaker 2: A 10 million core processor system for brain simulation and machine learning. arXiv preprint 1911.02385.

Meyer, A. C., Neher, E., and Schneggenburger, R. (2001). Estimation of quantal size and number of functional active zones at the calyx of held synapse by nonstationary epsc variance analysis. Journal of Neuroscience, 21(20):7889–7900.

Millidge, B., Tschantz, A., and Buckley, C. L. (2020). Predictive coding approximates backprop along arbitrary computation graphs. arXiv preprint 2006.04182.

Mnih, A. and Gregor, K. (2014). Neural variational inference and learning in belief networks. arXiv preprint 1402.0030.

Neal, R. and Hinton, G. (1998). A view of the em algorithm that justifies incremental sparse, and other variants. Learning in Graphical Models, pages 355–368.

Neftci, E. O., Pedroni, B. U., Joshi, S., Al-Shedivat, M., and Cauwenberghs, G. (2016). Stochastic synapses enable efficient brain-inspired learning machines. Frontiers in neuroscience, 10:241.

Oertner, T. G., Sabatini, B. L., Nimchinsky, E. A., and Svoboda, K. (2002). Facilitation at single synapses probed with optical quantal analysis. Nature neuroscience, 5(7):657–664.

Palacios, E. R., Isomura, T., Parr, T., and Friston, K. (2019). The emergence of synchrony in networks of mutually inferring neurons. Scientific reports, 9(1):1–14.

Pecevski, D., Kappel, D., and Jonke, Z. (2014). NEVESIM: Event-driven neural simulation framework with a python interface. Frontiers in Neuroinformatics, 8:70.

Peyser, A., Deepu, R., Mitchell, J., Appukuttan, S., Schumann, T., Eppler, J. M., Kappel, D., Hahne, J., Zajzon, B., Kitayama, I., et al. (2017). Nest 2.14. 0. Technical report, Jülich Supercomputing Center.

Pfister, J.-P. and Gerstner, W. (2006a). Beyond pair-based stdp: A phenomenological rule for spike triplet and frequency effects. In Advances in neural information processing systems, pages 1081–1088.

Pfister, J.-P. and Gerstner, W. (2006b). Triplets of spikes in a model of spike timing-dependent plasticity. Journal of Neuroscience, 26(38):9673–9682.

Pulido, C. and Ryan, T. A. (2020). Synaptic vesicle pools are a major hidden resting metabolic burden of nerve terminals. bioRxiv.

Ramstead, M. J., Veissière, S. P., and Kirmayer, L. J. (2016). Cultural affordances: Scaffolding local worlds through shared intentionality and regimes of attention. Frontiers in psychology, 7:1090.

Ramstead, M. J. D., Badcock, P. B., and Friston, K. J. (2018). Answering schrödinger’s question: A free-energy formulation. Physics of life reviews, 24:1–16.

Rao, R. P. and Ballard, D. H. (1999). Predictive coding in the visual cortex: a functional interpretation of some extra-classical receptive-field effects. Nature neuroscience, 2(1):79.

Rezende, D. J. and Gerstner, W. (2014). Stochastic variational learning in recurrent spiking networks. Frontiers in Computational Neuroscience, 8(38):1–14.

Rezende, D. J., Mohamed, S., and Wierstra, D. (2014). Stochastic backpropagation and approximate inference in deep generative models. In International conference on machine learning, pages 1278–1286. PMLR.

Rusakov, D. A., Savtchenko, L. P., and Latham, P. E. (2020). Noisy synaptic conductance: bug or a feature? Trends in Neurosciences, 43(6):363–372.

Schug, S., Benzing, F., and Steger, A. (2021). Presynaptic stochasticity improves energy efficiency and helps alleviate the stability-plasticity dilemma. Elife, 10:e69884.

Szavits-Nossan, J. and Evans, M. R. (2015). Inequivalence of nonequilibrium path ensembles: the example of stochastic bridges. Journal of Statistical Mechanics: Theory and Experiment, 2015(12):P12008.

Toyoizumi, T., Pfister, J.-P., Aihara, K., and Gerstner, W. (2005). Generalized bienenstock–cooper–munro rule for spiking neurons that maximizes information transmission. Proceedings of the National Academy of Sciences, 102(14):5239–5244.

Urbanczik, R. and Senn, W. (2014). Learning by the dendritic prediction of somatic spiking. Neuron, 81(3):521–528.

van De Burgt, Y., Melianas, A., Keene, S. T., Malliaras, G., and Salleo, A. (2018). Organic electronics for neuromorphic computing. Nature Electronics, 1(7):386–397.

Van Rossum, M. C., Bi, G. Q., and Turrigiano, G. G. (2000). Stable hebbian learning from spike timing-dependent plasticity. The Journal of Neuroscience, 20(23):8812–8821.

Whittington, J. C. and Bogacz, R. (2017). An approximation of the error backpropagation algorithm in a predictive coding network with local hebbian synaptic plasticity. Neural computation, 29(5):1229–1262.

Yang, H. and Xu-Friedman, M. A. (2013). Stochastic properties of neurotransmitter release expand the dynamic range of synapses. Journal of Neuroscience, 33(36):14406–14416.

Yger, P. and Gilson, M. (2015). Models of metaplasticity: a review of concepts. Frontiers in computational neuroscience, 9:138.

